# DyScore: A Boosting Scoring Method with Dynamic Properties for Identifying True Binders and Non-binders in Structure-based Drug Discovery

**DOI:** 10.1101/2021.10.26.465921

**Authors:** Yanjun Li, Daohong Zhou, Guangrong Zheng, Xiaolin Li, Dapeng Wu, Yaxia Yuan

## Abstract

The accurate prediction of protein-ligand binding affinity is critical for the success of computer-aided drug discovery. However, the accuracy of current scoring functions is usually unsatisfactory due to their rough approximation or sometimes even omittance of many factors involved in protein-ligand binding. For instance, the intrinsic dynamic of the protein-ligand binding state is usually disregarded in scoring function because these rapid binding affinity prediction approaches are only based on a representative complex structure of the protein and ligand in the binding state. That is, the dynamic protein-ligand binding complex ensembles are simplified as a static snapshot in calculation. In this study, two novel features were proposed for characterizing the dynamic properties of protein-ligand binding based on the static structure of the complex, which is expected to be a valuable complement to the current scoring functions. The two features demonstrate the geometry-shape matching between a protein and a ligand as well as the dynamic stability of protein-ligand binding. We further combined these two novel features with several classical scoring functions to develop a binary classification model called DyScore that uses the Extreme Gradient Boosting algorithm to classify compound poses as binders or non-binders. We have found that DyScore achieves state-of-the-art performance in distinguishing active and decoy ligands on both enhanced DUD dataset and external test sets with both proposed novel features showing significant contributions to the improved performance. Especially, DyScore exhibits superior performance on early recognition, a crucial requirement for success in virtual screening and *de novo* drug design. The standalone version of DyScore and Dyscore-MF are freely available to all at: https://github.com/YanjunLi-CS/dyscore

**Key Points:**

- Two novel binding features were proposed for characterizing the dynamic properties of protein-ligand binding only based on a static snapshot of complex.
- Based on the XGBoost machine learning method, the DyScore recognition model was proposed to accurately classify compound binding poses as binders or non-binders. DyScore consistently outperforms all the state-of-the-art published models on three different metrics by a large margin.
- DyScore showed superior performance in early recognition with an average of 73.3% success rate for the top three ranked compounds for each protein target.
- The standalone version of DyScore and DyScore-MF are freely available to all at: https://github.com/YanjunLi-CS/dyscore

**TOC:** 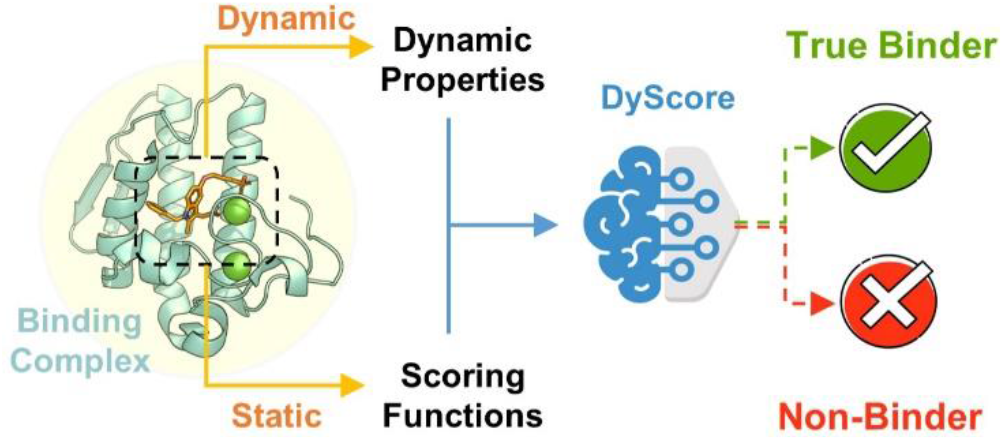

## 1 Introduction

Early-stage drug discovery focuses on identifying novel active compounds regulating drug targets specifically. The virtual screening (VS) method is widely used in computer-aided drug discovery in this stage^1^. However, its performance highly relies on the scoring function used for estimating the protein-ligand binding affinity. Despite notable improvements in accuracy and efficiency in past decades^2–12^, computational approaches still face considerable drawbacks and limitations. The protein-ligand binding affinity predicted by scoring functions usually suffer from significant biases with experimental data. As a result, false-positive predictions frequently occur in the practical drug discovery process, which is one of the major bottlenecks of computer-aided drug discovery method. Although there are many more precise computational techniques for protein-ligand affinity prediction, such as molecular mechanics/Poisson–Boltzmann or generalized Born surface area (MM/PBSA, MM/GBSA), linear interaction energy(LIE), thermodynamic integration (TI), free energy perturbation (FEP), and potential of mean force (PMF)^13–18^, these approaches are highly calculation-intensive and unrealistic for use in virtual screening. Additionally, it is also challenging to improve the prediction accuracy of scoring functions further because the partial absence of information about protein-ligand binding is an inherent trade-off between prediction accuracy and computational cost of scoring functions. The most apparent information missed by scoring functions is the dynamic properties of the protein-ligand binding process, which plays a crucial role in forming and stabilizing the binding complex. This is mostly because classical scoring functions are trained or calibrated based on protein-ligand binding complex structures solved by experimental approaches such as X-ray, NMR or Cryo-EM, which only provide one or a few static snapshots of the system in low energy state. However, although the static snapshot of complex structure does not explicitly present the mobility of ligand in the binding site, the potential fluctuation of ligand should be relevant to its interactions with protein residues. So, it would be possible to estimate the dynamic properties of protein-ligand binding based on the static snapshot of a complex structure.

In this study, two novel features were proposed for characterizing the dynamic properties of protein-ligand binding based on the static structure of a complex, and they are expected to be a valuable complement for current scoring functions. It is worth noting that the two features related to dynamic properties are distinct from typical energy-based terms used in classic scoring functions. For example, the ligand flexibility terms are widely used in scoring functions, as an empirical estimation of conformational entropy difference between the bound and unbound state of ligand. It is a part of the schema of calculating Gibbs free energy. In this schema, protein flexibility is more important for entropy change. However, due to the complexity of protein structure, although many MD simulation-based and normal-mode-based approaches were developed to calculate the entropy difference of protein in the bound and unbound state, it is still very challenging to estimate the conformation entropy in either aspect of time consumption or accuracy. So, our principal idea is to look out of the Gibbs free energy schema which is established at the equilibrium state. We tried to estimate the dynamic properties of the binding process. As Gibbs free energy would be the theoretical basis of complex-ligand binding, any estimation of binding properties from other aspects should have to partially overlap with Gibbs free energy terms, including the entropy penalty from ligand flexibility terms and protein flexibility terms. We expect to use a non-Gibbs free energy schema to reveal more information in the binding process that could not be accurately estimated in the Gibbs free energy schema. The two novel features are essentially different from the typical energy-based terms in the Gibbs free energy schema. That is, they could not be directly decomposed into individual energy terms contributing to binding free energy because they demonstrate a comprehensive favorability of binding, which involved various entropy and enthalpy items as well as entropy-enthalpy coupling. As a result, the two features were not simple to integrate into classic scoring functions because they partially overlap with other features in scoring function. Therefore, the machine learning method was used in this study to comprehensively combine scoring functions with the two new features. The training objective of the machine learning model could be either binding affinity prediction or binder/non-binder recognition. Although the binder/non-binder recognition is a subset problem of binding affinity prediction, the model designed for binding affinity prediction usually shows significant performance degradation in binder/non-binder recognition^11, 19^. This bias is principally rooted in the type of training data used to construct the model. For a binding affinity prediction model, training data is usually composed of only active compounds with known binding affinity. This results in the model implicitly assuming that all input data are active and easily being misled by decoy ligand in practice. On the contrary, a binder/non-binder recognition model learns to identify active compounds from massive decoys, which approximates more to the common scenario of virtual screening and *de novo* design, therefore make it more important for practical applications. In this study, a novel binder/non-binder recognition model named DyScore was constructed to identify true binders and non-binders for structure-based drug discovery. Within our knowledge, this is the first work that incorporates the static structure and its dynamic properties of the protein-ligand complex in scoring function to improve the binding affinity prediction. Extensively evaluated using the commonly used benchmark dataset DUD-E and an external test set DEKOIS 2.0, DyScore was found to consistently outperform all the state-of-the-art (SOTA) published models on three different metrics by a large margin, especially within early recognition, which is a crucial requirement for success in virtual screening and *de novo* drug design. We demonstrated the notable feature importance of the two proposed dynamic properties for classification performance in the interaction-based machine learning model.

## 2 Materials and Methods

### 2.1 Dataset Preparation

#### Data provenance

The Directory of Useful Decoys-Enhanced (DUD-E)^20^ dataset is broadly used as a recognized benchmark for estimating the performance of VS algorithms and scoring functions. The dataset contains 102 protein span diverse protein categories, including 26 kinases, 15 proteases, 11 nuclear receptors, 5 GPCRs, 2 ion channels, 2 cytochrome *P450s*, 36 other enzymes, and 5 miscellaneous proteins, which is a representative set for typical drug targets. A total 22, 886 known active ligands are drawn from ChEMBL^21^ (average 224 active ligands for each target), each with average 61 property-matched decoys drawn from ZINC database^22^ (average 61 decoys per active ligand). All the active ligands in the DUD-E are reported to bind with corresponding protein targets with a binding affinity (*IC*_50_, *EC*_50_, *Ki*, or *K_d_*) better than 1*μ*M. In contrast, the decoy ligands are selected to mimic the distribution of physical properties (including molecular weight, logP, number of rotatable bonds, and hydrogen bond donors and acceptors) of active ligands on a target-by-target basis. The topological similarity between decoys and actives is minimized by using 2D similarity fingerprints. Therefore, the decoys in the DUD-E dataset are expected to resemble the physical properties of active ligand but be topologically dissimilar to minimize the likelihood of actual binding, which would be challenging for docking algorithms and scoring functions. In this study, the DUD-E dataset was used to generate training, validation, and test sets for constructing and evaluating our method.

To externally evaluate our approach, we also used DEKOIS 2.0^23^ and LIT-PCBA^24^ dataset as independent test sets. The DEKOIS 2.0 library includes 81 benchmark sets for 80 protein targets (2 benchmark sets for different PYGL binding sites), and it comprises a diverse selection of protein classes like kinases, proteases, nuclear receptors, GPCRs, oxido-reductases, transferases, hydrolases, and several others. Because 4 out of 81 targets of the DEKOIS are also in the DUD-E data set, we removed 4 targets (A2A, HDAC2, PARP-1, and PPARA) from the DEKOIS set to avoid data leakage and used the remaining 77 targets to evaluate our model’s performance. For simplicity, in the rest part of the article, we consistently use DEKOIS 2.0 data set to represent the filtered DEKOIS set, which contains 77 targets and no-shared targets with the DUDE data set. The LIT-PCBA dataset consists of 15 targets and 407,381 confirmed inactive compounds and 7844 confirmed active compounds. The LIT-PCBA dataset is developed to mimics the hit rate (ratio of active to inactive compounds) and potency distribution in experimental decks. We did not remove duplicate target from LIT-PCBA for comparison of results with other studies that also kept the duplicates.

#### Generation of ligand binding poses

The initial 3D conformations of active and decoy ligands in the DUD-E, DEKOIS 2.0 and LIT-PCBA datasets were generated by the OMEGA^25^ module of OpenEye software. Then Fixpka^26^ module of OpenEye software was used to set the correct protonation status for each compound at *pH* = 7. Autodock Vina (Vina)^27^ was used to dock all the active and decoy ligand in the DUD-E dataset to its corresponding protein target. GNU parallel is used for parallel processing^28^. From the total 20 protein-ligand conformations generated by Vina, the conformation with the lowest binding energy was selected for further analysis. The binding site box of each protein target for conformation searching in protein-ligand docking was defined based on the ligand in the corresponding crystal structure. The box center was aligned to the mass center of the ligand, and its size was set to fitting the ligand with an extended boundary of 10Å.

#### Data split

In order to follow the practical application scenarios of the scoring functions and comprehensively evaluate our proposed method, we introduced two different data split paradigms, named target-aware split and target-unaware split. **The target-aware split** represents the scenario when computing the binding score of a novel ligand to a target protein, we have already known some other active ligands of this target. As shown in **Figure 1a**, the binding complexes for each target were randomly split into three non-overlapped subsets with the ratio of 80% - 10% - 10% on a target-by-target basis, and both active and decoy data were split separately in the same manner to maintain the consistent distribution of active and decoys in subsets for each target. Then the corresponding subsets across 102 targets were merged to construct the training, validation, and test sets, respectively. In total, the training set contains 1,139,271 complexes, where the numbers of actives and decoys are 18,197 and 1,121,074. The complexes in either of the validation and test sets are around 1/8 of the training set. As the diversity, pharmacological precedence, target relevance, and ease of study were considered in target selection of DUD-E set^20^, the frequency of occurrence and type of targets in DUD-E set represents a typical expectation for new drug targets. So, this splitting approach was used to preserves the data distribution of different targets in the final three subsets as same as the original DUD-E set.

**Fig. 1:**
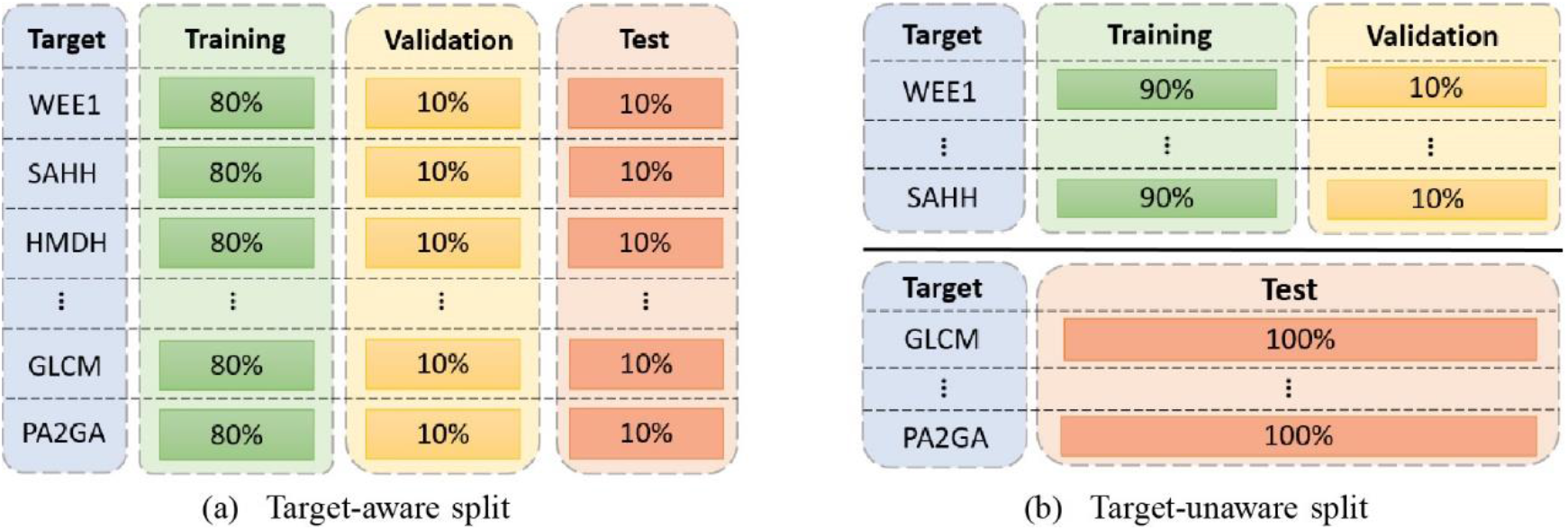
Illustration of the data split on the DUD-E data set. Note that different targets may contain different numbers of complexes, although we use the horizontal bars with the same length to represent.

The **target-unaware split** aims to represent the scenario in which the target is unseen to the scoring function, and it does not have any known ligand. As illustrated in **Figure 1b**, we first randomly selected 11 targets (around 10% out of 102 targets in total), and merged all the binding complexes of these targets as the test set. For the remaining complexes, we randomly split them into two non-overlapped training and validation subsets with the ratio of 90% - 10% on a target-by-target basis. In order to make the equivalent distribution of active and decoy complexes in both subsets for each target, both active and decoy data were split independently with the ratio of 90% - 10%. In summary, the training and validation sets have no shared target with the test subset under the target-unaware split.

For the above two data split manners, all random splits were independently repeated three times to generate three parallel data sets, and the models were trained and evaluated on these three parallel data sets to investigate their robustness. The mean performance of three models was reported as the final performance.

### 2.2 Generation of Lock-key Matching Score Feature

According to the classical lock-key model proposed by Emil Fischer in 1894^29^, a typical protein-ligand binding complex usually achieves high geometric complementarity, meaning the spatial fitting of the geometric shape of a ligand to a binding site is favorable for typical protein-ligand binding. In this study, a lock-key matching score was proposed to characterize the geometric complementarity between proteins and ligands. The definition of our proposed lock-key matching score is depicted in **Figure 2.** It would be intuitive to demonstrate the geometric complementarity by evaluating the degree of motion freedom for ligand in the ligand binding site. One common strategy is to calculate the ratio of the region occupied by protein and unoccupied region (vacant region) around ligand; however, this ratio does not consider the size of the ligand, which is necessary for evaluating geometric complementarity. To address this, our proposed lock-key matching score is determined by analyzing the motion accessible and inaccessible region of the ligand in the binding site, specifically, by calculating the volume difference between occupied and vacant regions around the ligand, which is expected to reflect the extent of geometric complementarity. To calculate the volume of the irregular geometric shape of protein and ligand, 3-dimensional (3D) lattice space is used to split the occupied and vacant regions into small countable cubic grids. So, the total volume of each type of region could be estimated by the total number of corresponding grids. Considering the van der Waals radius of common heavy elements in protein and ligand are in the range of 1.5Å to 1.8Å the edge length of the cubic grid is defined as 0.5Å which is sufficient to characterize the shape of atom surface. We do not adopt a smaller edge length such as 0.25Å or 0.1Å because smaller edge length will also characterize the small interstices in van der Waals surface of molecules as vacant region, which is not accessible by any atoms and therefore should not be considered as a vacant region. Besides, the smaller edge length than 0.5Å also increases the fluctuation and noise of results due to its sensitivity to tiny conformational change.

**Fig. 2:**
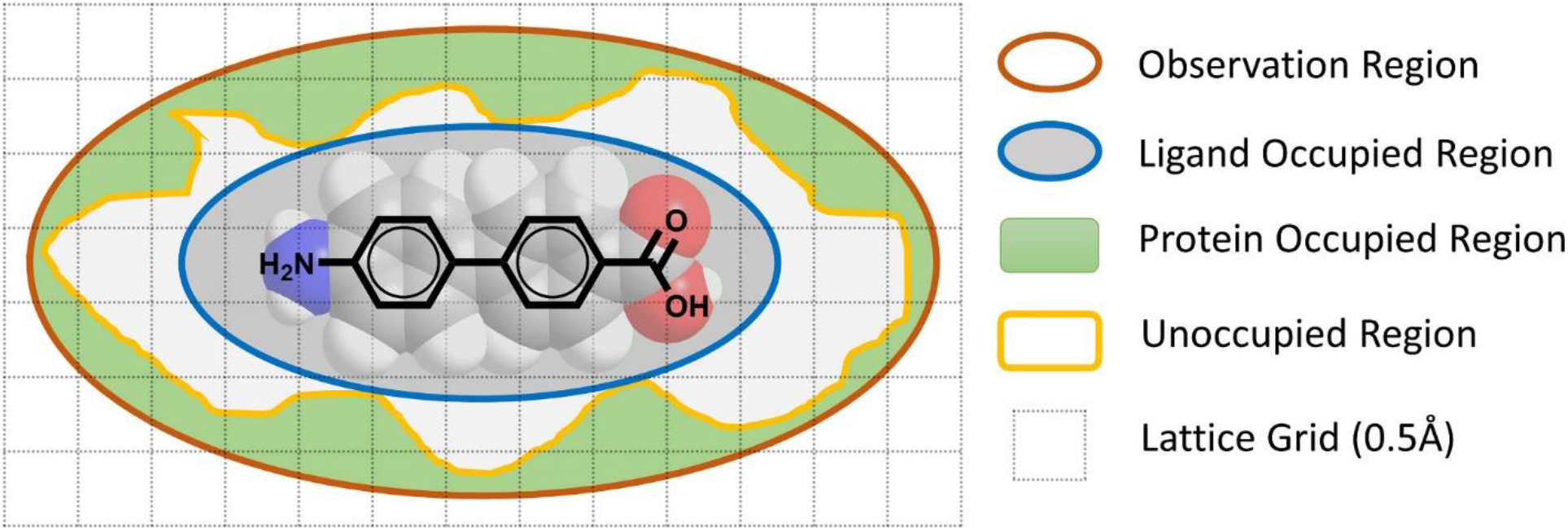
Definition of the lock-key matching score. The region colored in green indicates space occupied by atoms of the protein. The white region indicates space that is not occupied by atoms of protein or ligand. The gray region in the blue cycle indicates the space occupied by atoms of ligand. The matching score is defined by the difference between the green and white regions. (Only grids within the observation region are accounted in.)

In the first step, residues around the ligand binding site of the target protein are identified by using a binding site detection program Cavity^30–32^. Then residues involved in the binding site as well as the ligand are mapped to the 3D lattice space. If the center of a cubic grid is located within the van der Waals radius of an atom from protein residues or ligand, this grid will be defined as an occupied grid; Otherwise, the grid is defined as a vacant grid. Specifically, we use *P_grid_, L_grid_*, and *V_grid_* to represent the grid occupied by atoms of protein residues, atoms of ligand, and unoccupied region (vacant region), respectively. Then the lock-key matching score is calculated by the number of *P_grid_* minus the number of *V_grid_*. It worth noting that the matching score, in essence, estimates the local gaps between protein and ligand, so only *P_grid_* and *V_grid_* nearby the ligand is calculated. Considering the diameter of the water molecule is about 2.8Å adding to the van der Waals radius of a heavy atom in the range of 1.5Å to 1.8Å the maximal observation region is defined by the grids around the ligand within 4Å to cap off the contribution of water accessible region gaps and solvent exposed grids.

### 2.3 Generation of Protein-Ligand Binding Stability Score Feature

As a result of the random thermal motion of protein, ligand, and solvent molecules, the protein-ligand interaction is actually a dynamic concept instead of a static scene. The ligand in a protein-ligand complex is constantly perturbed by its thermal fluctuation and random hitting from solvent molecules, which eventually induces the disassociation of ligand from the complex. Based on the classic protein-ligand binding theory, if a ligand could stably stay in the binding site, it is likely to achieve higher binding affinity due to its slower disassociation process (smaller disassociation constant *K_off_*). Based on this principle, a mechanical model was proposed to estimate the stability of ligand binding to a protein.

Because the calculation in this study is based on a static snapshot of the protein-ligand complex and the complex’s dynamic information is missing, the stability of the ligand is estimated by its degree of freedom. The definition of protein-ligand binding stability score is depicted in **Figure 3**. Here we defined two types of potential force which could be restrictions on the degree of freedom of ligand. One is the potential repulsion force (source from van der Waals interaction), which could provide a counterforce if the ligand is closer to the protein atom than the equilibrium distance. The other one is potential attraction force (source from hydrogen bond, metal coordination, or electrostatic interaction), which could provide a counterforce to prevent the ligand from leaving. The distance cutoff of van der Waals contact between two atoms is defined with the sum of their van der Waals radii plus 0.5Å. The definition of polar interactions follows the methods described in Score2^33^. Although the net force of all potential forces is negligible in an equilibrium complex, the potential forces are expected to constrain both translational and rotational degrees of freedom of the ligand. Based on this principle, we calculate the translational stability and rotational stability of ligand by estimating the losing of translational and rotational degrees of freedom under the restriction of potential forces. In consideration of the possible internal flexibility of a molecule, some ligands may not be treated as an intact rigid body in calculating the degree of freedom. In this case, the ligand is split into multiple rigid fragments by cutting all non-ring flexible bonds, and the stability of each rigid fragment will be calculated independently (**Figure 3b**). However, the flexible bonds between rigid fragments also restrict the degree of freedom of each fragment, so it is treated as a pair of bidirectional virtual attraction and repulsion force in the calculation, which preserves the force conduction from its neighbor fragments (**Figure 3c**). Then the loss of a degree of freedom for a rigid fragment is calculated by summing up the restriction of all potential forces on the rigid fragment. It should be noted that the calculation is based on local perturbation of ligand instead of the actual dissociation process. That is, all the protein-ligand interactions in the complex are not broken in calculating the dynamic features of the ligand. So, the strength of all the potential forces is not considered; instead, the direction of the potential force on the rigid body is more important for estimating the short-range restriction of the degree of freedom.

**Fig. 3:**
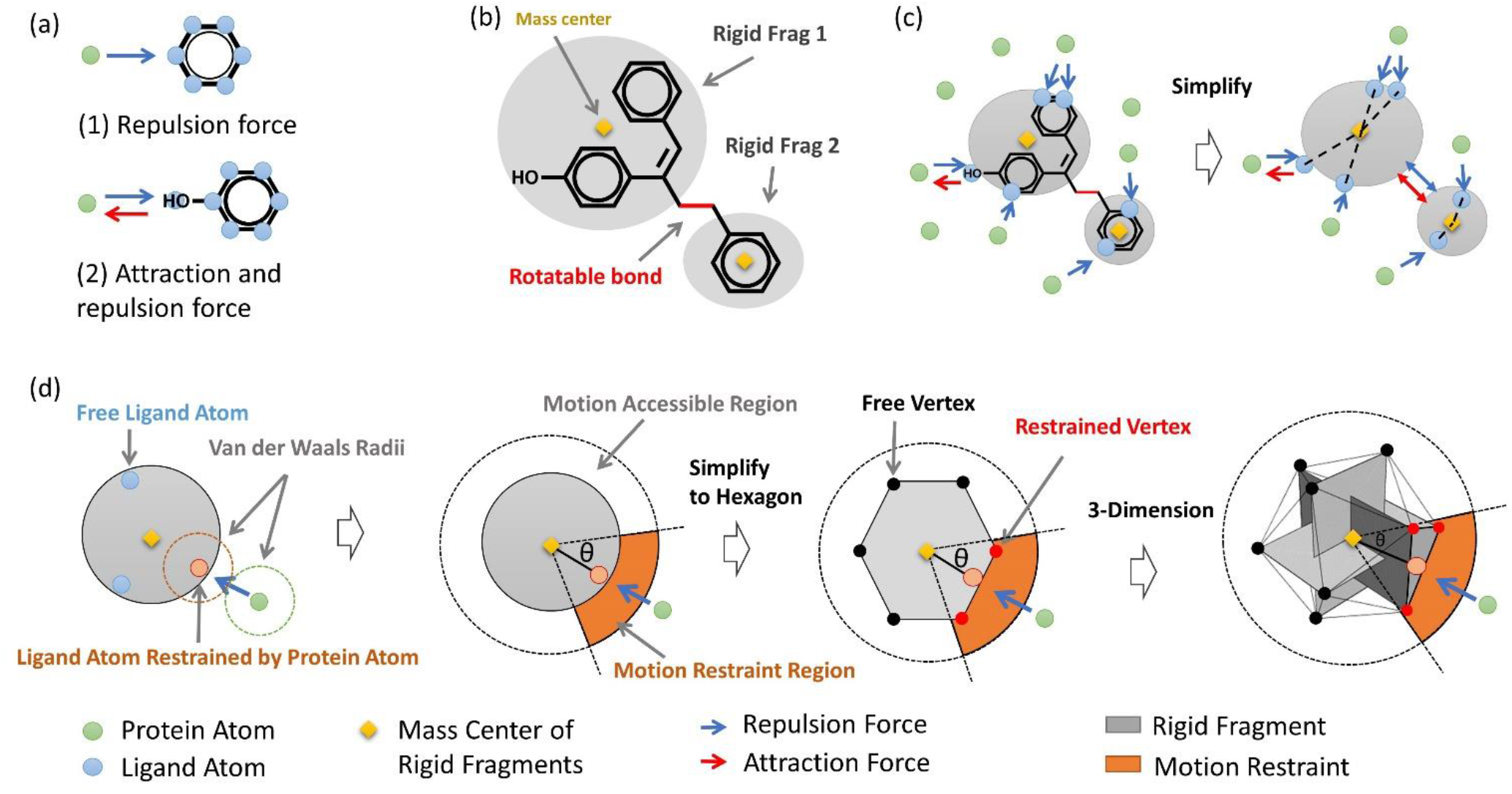
(a) Illustration for two types of potential force between protein and ligand. One is the potential repulsion force (source from van der Waals interaction), which could provide a counterforce if the ligand is closer to the protein atom than the equilibrium distance. The other one is potential attraction force (source from hydrogen bond, metal coordination, or electrostatic interaction), which also could provide a counterforce to prevent the ligand from leaving. The potential repulsion force and attraction force are represented by blue and red arrows, respectively. Potential repulsion force is always coupling with possible attraction force. (b) the rigid fragments split by rotatable bonds in ligand could be treated as several separated rigid bodies. (c) The comprehensive effect of all potential repulsion and/or attraction forces for each rigid body could be considered as a resultant force. Intra-force between rigid body is treated as bidirectional repulsion and attraction force. (d) The potential repulsion and/or attraction force between protein and ligand would result in motion restraints of ligand, which is intuitive to demonstrate in the two-dimension model. By simplifying the rigid body into a hexagon in a 2D model, the motion restraint could be evaluated by counting the restrained vertex (colored in red) and free vertex (colored in black). By using icosahedron (12 vertexes) to demonstrate the rigid body, the motion restraint originated from protein-ligand interaction could be evaluated in the same manner as the two-dimension model.

As depicted in **Figure 3d**, a regular icosahedron model with 12 vertexes is proposed to approximately demonstrate the degree of freedom of ligand by the number of free vertexes of the icosahedron. Subsequently, the loss of a degree of freedom of ligand, namely, stability of ligand, is approximately defined as the ratio of constrained vertexes among all 12 vertexes. Take the translational degree of freedom as an example. Any translational movement in 3D space could be decomposed into three orthogonal movement vectors along the X, Y, Z-axis. So there is a total of 6 uncoupled movement directions, that is, X, -X, Y, -Y, Z, -Z, which suggests that at least 6 vertexes are required for using a regular polygon model to demonstrate the degree of freedom of an object. Although it would be simplest to adopt a regular octahedron model with 6 vertexes, the 6 vertexes are uncoupled to each other, which would be more sensitive to small conformational perturbation if the movement is coincidentally around the 45° boundary between two orthogonal axes, so we adopt a regular icosahedron model with 12 vertexes instead to increase the robustness of calculation.

The translational stability of each rigid fragment is calculated using the following steps: **(a)** The rigid fragment is simplified as a regular icosahedron with 12 vertexes, and the center of the icosahedron is aligned to the mass center of the rigid fragment. For each vertex, we define a vertex vector from the center of the icosahedron to this vertex. **(b)** The repulsive forces between an atom of protein and the related rigid fragment of ligand will be simplified to force vectors from the ligand atom pointing towards its contacted protein atom. The attractive forces are simplified to force vectors from protein atoms pointing towards their contacted ligand atoms. **(c)** If the vectorial angle between a force vector of either repulsive or attractive force and any vertex vector is less than 32°, the corresponding vertex is labeled as a stabilized vertex. Each stabilized vertex indicates a direction that the rigid fragment will be block from moving towards. As the vertex-center-vertex angle of the icosahedron is about 63°, the 32° threshold will guarantee each force vector could be allocated to at least one vertex. **(d)** The total translational stability of the rigid fragment is then calculated by dividing the number of stabilized vertexes by 12, the total number of vertexes of an icosahedron.

The rotational stability of each rigid fragment is calculated using the following steps: **(a)** Define the vertex vector. **(b)** Define the force vector. Both these steps use the same calculation methods as used to determine translational stability. **(c)** A moment vector is derived for each force vector using the dot product of the force vector and the vector from the center of icosahedron pointing to the force vector, which indicates the potential effect of force on fragment rotation. **(d)** If the vectorial angle between a moment vector of either repulsive or attractive force and any vertex vector is less than 32°, the corresponding vertex is labeled as a stabilized vertex. Each stabilized vertex indicates a direction that the rigid fragment will be blocking from rotating towards. **(e)** The total rotational stability of the rigid fragment is then calculated by dividing the number of stabilized vertexes by 12, the total number of vertexes of an icosahedron.

Finally, the translational stability and rotational stability of ligand are defined by the mean stability of each rigid fragment. The calculated translational and rotational stability could be considered as an approximation of the average stability of 3 translational and 3 rotational degrees of freedom for the ligand, respectively. As the stability of all 6 degrees of freedom are related to the dissociation of ligand from protein, the comprehensive stability of ligand is defined by the product of stability in 6 degrees of freedom; that is, the third power of average translational stability multiplied by the third power of average rotational stability.

### 2.4 Generation of Other Features for DyScore

In addition to the two proposed novel features, we have developed for determining the dynamic properties of protein-ligand binding, several classical scoring function outcomes and several basic static binding features were included as inputs for DyScore. This included three categories of classic scoring functions for binding affinity estimation based on static complex structures: (1) empirical scoring function (Vina Score^27^, Score2^33^, and X-Score^34^); (2) pairwise potential based scoring function (DSX^35^); (3) machine learning scoring function (RF-Score V3^36^, and NNscore V2^37^). To avoid introducing underlying bias from a scoring function embedded with information from the DUD/DUD-E dataset, we only selected scoring functions that were not trained on the DUD/DUD-E dataset. Additionally, five basic static binding features related to protein-ligand binding (calculated by X-Score^34^) were used as input features, including van der Waals energy, count of metal coordinate bonds, count of hydrogen bods, hydrophobic matching for protein-ligand bonding, and the number of rotatable ligand bonds. The distribution of each input feature in DUD -E data set and DEKOIS 2.0 data set can be found in Supplementary Figure S1 and S2. It is worth noting that as a general framework, DyScore can integrate some additional powerful scoring functions such as DeepAtom^38^, KDeep^39^, and RF-Score^40^. However, a thorough integration of these scoring functions is beyond the scope of this work, so we plan to explore the benefits of combining more binding scores and features in future work.

### 2.5 Generation of Molecular Fingerprints (MF) for DyScore-MF

DyScore-MF model is trained based on all input features used in DyScore model, with an extra FP2 molecular fingerprint (MF) generated by OpenBabel^41^. FP2 is a widely used path-based fingerprint which indexes small molecule fragments based on linear segments of up to 7 atoms. It encodes both fragments and local topological information of target molecules via a 1024 binary bit vector. Combined with 14 input features used in DyScore, the total number of final input features for DyScore-MF is 1038. It encodes both fragments and local topological information of target molecules. There are 14 input features used in DyScore, and the FP2 fingerprint is represented by a 1024 binary bit vector, so the total number of combined input features for DyScore-MF is 1038.

### 2.6 Machine Learning Algorithm of DyScore and DyScore-MF

The Extreme Gradient Boosting (XGBoost) was used as the learning algorithm of DyScore. XGBoost is an ensemble machine learning method that applies the boosting strategy to combine multiple base learners into a powerful and robust model^42^. Evolved from the gradient tree boosting algorithms, XGBoost is designed as a highly scalable end-to-end system with improved computational efficiency and a more potent capacity to alleviate the overfitting problem. It has achieved SOTA results on a broad range of machine learning tasks, especially learning from the structured or tabular datasets with highly diverse and complex feature space and imbalanced class distribution.

For the XGBoost algorithm of DyScore, the random forest was selected as the homogeneous base learner for gradient boosting. Although some weak machine learning models (e.g., decision trees) are commonly used as the base learner, we observed the noticeable performance improvement by employing random forest in the DyScore XGboost, which was further compared in Section 3.4. The binary logistic regression was set as the training objective, and several strategies were utilized to prevent overfitting. Specifically, the boost learning step was slowed down by setting the shrinkage factor to 0.05, and the early stopping regularization was used to terminate the training process if the average per-target Area Under the Curve (AUC) score on the validation set did not improve in the consecutive 20 boost rounds. The weight for L2 regularization was set as 1 to reduce the model complexity. For RF-specific parameters, the subsample ratio was set as 0.85 to randomly select 85% training samples before adding a new RF in every boosting iteration and subsample 30% features when splitting for each node. Additionally, to reduce the class-imbalance impact introduced by the data distribution of the DUD-E dataset, the scale for positive weight was set as 60, which can facilitate the model controlling the balance of the positive and negative samples’ weights.

### 2.7 Evaluation Metrics

In this study, several broadly used metrics were applied to evaluate the model comprehensively from different aspects. The AUC of Receiver Operating Characteristic (ROC)^43^ measurement was used for the classification performance assessment. It can effectively demonstrate the global classification performance throughout the ranked prediction list. The model’s performance on early recognition was also an important aspect to evaluate because the accuracy of true binder recognition within the model’s top predictions plays a crucial role in the success of practical VS and *de novo* drug design. Therefore, the Enrichment Factor (EF) (shown in the ***Eq. 1***) was used to measure how much the presence of the active molecules within a specific top range (or percentage) of the ranked predictions has increased relative to a random selection:

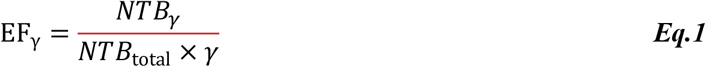

*NTB_γ_* is the number of the true binders among the top-ranked predictions with different early recognition cutoff *γ* settings (e.g., 1%, 2%). *NTB*total is the number of true binders for a target protein in the total prediction list. Note that the ratio of randomly selecting an active molecule out from the DUD-E dataset is around 1.6% because the positive/negative ratio is 1:61 based on the distribution of the actives and decoys. For the DEKOIS 2.0 dataset, the ratio is 1:30. Although EF can evaluate the model capacity purely based on the top-ranked predictions, it still has a limitation that it equally weights all the actives falling in this range. Therefore, the Boltzmann-Enhanced Discrimination of the Receiver Operating Characteristic (BEDROC)^44^ (***Eq. 2***) was additionally employed as the other early recognition metric, which can adaptively weight the actives based on their ranking positions. Suppose a ranked prediction list contains *N* molecules, and the number of actives is *n*. For *i*th active molecule, *r_i_* represents its ranking position in the full list and *x_i_* is its relative rank where *x_i_* = *r_i_*/*N*. The BEDROC is calculated as follows:

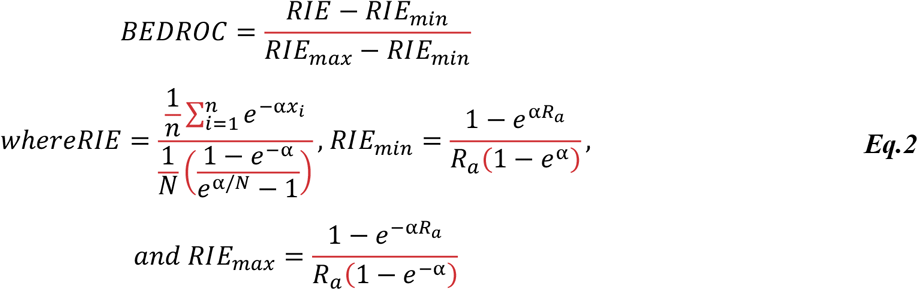

*RIE* is short for the Robust Initial Enhancement metric, *R_a_* = *n/N* and *α* is an early recognition parameter. For instance, with setting *α* = 160.9, 80% of the overall BEDROC score is contributed by the top 1% of the ranked predictions. Note that the prediction performance on a single metric varies from target to target. Following the conventional practice, the averaged results of 102 different targets were served as the overall performance of the model.

### 2.8 Methods for Comparison

On the one hand, DyScore was compared with six SOTA scoring methods: RF-Score-v3, NNscore-v2, X-Score, Score2, DSX, and Vina Score. We used the publicly available binaries provided by the authors of these compared methods to make predictions on the same validation and test sets. It is worth noting that these methods were exclusively trained on the protein-ligand complexes crystal structures in the PDBbind data set. Because of negligible overlap between ligands in PDBbind and DUD-E data sets, these methods were unlikely to introduce the similarity-based information discussed in Section 3.3.

On the other hand, four different machine learning and deep learning methods were evaluated based on the same 14 binding features. Besides the random forest based XGBoost model adopted by DyScore, another XGboost based method (DyScore-DT-XGboost) with decision tree as its base learner was developed for comparison. The training strategies of the boosting algorithm were kept the same with DyScore to prevent overfitting. We also compared a Random Forest model (DyScore-RF) with 500 parallel trees. In order to prevent overfitting and improve the tree diversity, the max depth of each tree was constrained as 40, randomly subsampled 4 features are considered for each node splitting, and the minimum number of samples was set as 5 for a node to become a leaf. In addition, we also developed a deep neural network based approach, termed as DyScore-MLP. MLP stands for multiple-layer perceptron model, a common network structure used to learn from structured/tabular data. 7 dense layers with 512 hidden units were sequentially stacked, and a batch normalization operation was also appended after each dense layer. ReLU was used as the nonlinear active function and the binary cross-entropy loss was used for optimization. For the DyScore-MLP model, all the input features were scaled to the same range of [1, 1] before feeding into the network. All competing methods use the same early stopping regularization as DyScore, where the training process terminated if the average per-target AUC score on the validation set did not improve in the consecutive 20 learning epochs, and hyper-parameters of all the methods were extensively optimized on the same validation set for a fair comparison.

## 3 Results and Discussion

The performance of DyScore with the two proposed features was systematically evaluated on the DUD-E benchmark and external DEKOIS 2.0 and LIT-PCBA dataset to demonstrate its capacity to identify true binders and non-binders. The ranking power of DyScore was compared with 6 SOTA scoring functions in Section 3.1 under the unified comparison metrics described in Section 2.6. In Section 3.2, the permutation importance method^45^ was used to demonstrate the significance of matching score and stability score by measuring how performance decreased when they were not available in the input features. DyScore is designed as an interaction-based model which focuses on improving the generalization ability, but we also discussed a similarity-based model DyScore-MF in Section 3.3. The performances of several machine learning and deep learning methods training on the same features were further assessed in Section 3.4. In Section 3.5, we compared the performances on several key early recognition metrics, which are crucial for *de novo* design.

### 3.1 Comparison of DyScore and Other SOTA Scoring Methods

In this section, DyScore was compared with the six SOTA scoring methods: RF-Score-v3, NNscore-v2, X-Score, Score2, DSX, and Vina Score. The evaluation was conducted over three different data settings: **(1)** target-aware split for DUD-E data set (see 3.1), **(2)** target-unaware split for DUD-E data set (see 3.1), **(3)** external DEKOIS 2.0 benchmark (see 3.1), and **(4)** external LIT-PCBA benchmark (see Supplementary Result 1.1). For the first two settings, all the methods were evaluated on the three independent random split test sets, and the final performance was reported by the average of metrics for three test sets.

#### Evaluate on the DUD-E test set with target-aware split

This experimental setting mimicked the scenario in which the target protein in the protein-ligand complex has some other known active ligands. The DyScore was developed based on the 90% of complexes (80% for training and 10% for validation) over all the 102 DUD-E targets and then tested on the remaining 10% complexes. The distribution and median of AUC, EF-1%, BEDROC (321.9), and EF-top3 on the test set were depicted in the box and whisker plot **Figure 4**, where BEDROC (321.9) corresponds the top 0.5% predictions contributes 80% to the overall BEDROC score. DyScore outperforms all the other scoring methods over all the evaluation metrics by a large margin, especially for the early recognition measurements. DyScore achieves the EF-top3 of 45.1, and BEDROC (321.9) of 0.644, which is significantly higher than the second-best method Vina score with 14.4 and 0.213, respectively. The relative improvements on EF-top3 and BEDROC (321.9) by DyScore are around 213.2% and 202.3%. The full comparison results are shown in the first part of the **Table 1,** where the best performance of each evaluation metric is bold. The comparison results on the validation set can be found in **Supplementary Table S1** and **Figure S3**, and 102 per-target ROC curves generated by DyScore are shown in **Supplementary Figure S7**.

**Fig. 4:**
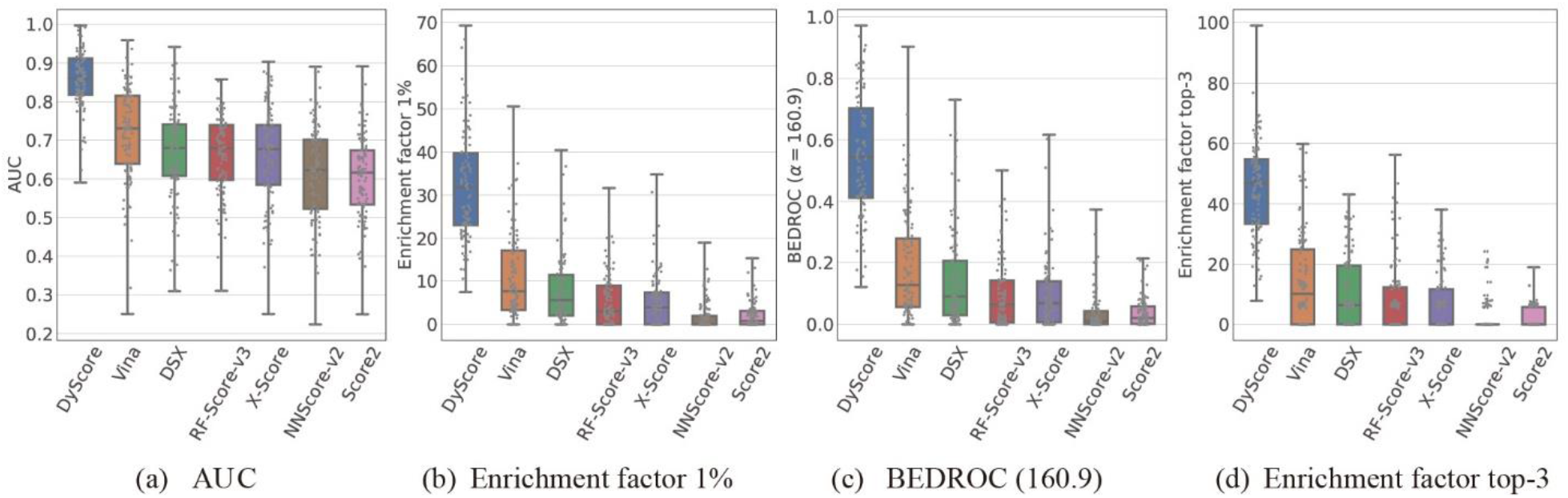
Comparison of different scoring methods on the DUD-E test set generated by target-aware split. Each gray dot represents a target (102 in total), and every point value is averaged over three randomly split test sets.

**Table 1:** Comparison of DyScore with six SOTA scoring functions and three other machine and deep methods learning from the same features. All the performances were evaluated on the DUD-E test set generated by the three randomly target-aware splits, the mean value was reported as well as the standard deviation in parentheses.

| Methods | AUC | Enrichment Factor |  |  |  |  |  | BEDROC |  |  |  |  |
| --- | --- | --- | --- | --- | --- | --- | --- | --- | --- | --- | --- | --- |
|  |  | Top 3 | Top 5 | 1% | 2% | 5% | 10% | 321.9 | 160.9 | 80.5 | 32.2 | 16.1 |
| **DyScore** | **0.858**<br>**(0.003)** | **45.1**<br>**(1.2)** | **40.1**<br>**(0.3)** | **32.2**<br>**(0.6)** | **21.2**<br>**(0.6)** | **10.9**<br>**(0.2)** | **6.4**<br>**(0.1)** | **0.644**<br>**(0.003)** | **0.551**<br>**(0.006)** | **0.502**<br>**(0.009)** | **0.514**<br>**(0.009)** | **0.564**<br>**(0.007)** |
| Comparison of DyScore and other SOTA Scoring Methods |  |  |  |  |  |  |  |  |  |  |  |  |
| Vina | 0.720<br>(0.008) | 14.4<br>(0.7) | 12.8<br>(0.8) | 10.8<br>(0.9) | 7.9<br>(0.7) | 5.3<br>(0.4) | 3.7<br>(0.2) | 0.213<br>(0.008) | 0.190<br>(0.012) | 0.187<br>(0.014) | 0.225<br>(0.015) | 0.288<br>(0.015) |
| DSX | 0.670<br>(0.006) | 10.9<br>(1.3) | 9.6<br>(0.7) | 8.0<br>(1.2) | 6.6<br>(0.7) | 4.3<br>(0.4) | 3.0<br>(0.1) | 0.163<br>(0.020) | 0.147<br>(0.017) | 0.149<br>(0.015) | 0.185<br>(0.014) | 0.238<br>(0.013) |
| RF-Score-v3 | 0.664<br>(0.007) | 7.9<br>(0.6) | 7.3<br>(0.3) | 5.5<br>(0.1) | 4.5<br>(0.2) | 3.3<br>(0.1) | 2.6<br>0.0 | 0.108<br>(0.002) | 0.099<br>(0.002) | 0.103<br>(0.004) | 0.138<br>(0.003) | 0.194<br>(0.002) |
| X-Score | 0.661<br>(0.008) | 6.9<br>(0.8) | 6.3<br>(0.5) | 5.2<br>(0.2) | 4.5<br>(0.3) | 3.4<br>(0.1) | 2.7<br>0.0 | 0.104<br>(0.005) | 0.097<br>(0.004) | 0.103<br>(0.005) | 0.140<br>(0.006) | 0.197<br>(0.006) |
| NNScore-v2 | 0.614<br>(0.004) | 2.3<br>(0.9) | 2.1<br>(0.7) | 1.7<br>(0.1) | 2.0<br>(0.1) | 1.9<br>0.0 | 1.8<br>0.0 | 0.033<br>(0.010) | 0.035<br>(0.006) | 0.044<br>(0.003) | 0.075<br>(0.003) | 0.126<br>(0.003) |
| Score2 | 0.607<br>(0.005) | 2.1<br>(0.5) | 2.3<br>(0.5) | 2.1<br>(0.2) | 2.1<br>(0.2) | 2.1<br>(0.1) | 1.9<br>0.0 | 0.034<br>(0.007) | 0.037<br>(0.006) | 0.048<br>(0.005) | 0.081<br>(0.005) | 0.132<br>(0.005) |
| Comparison of DyScore and other methods learning from the same features |  |  |  |  |  |  |  |  |  |  |  |  |
| DyScore-MLP | 0.838<br>(0.003) | 35.2<br>(1.3) | 31.9<br>(2.2) | 25.2<br>(2.2) | 17.6<br>(1.0) | 9.7<br>(0.2) | 6.0<br>(0.1) | 0.506<br>(0.029) | 0.441<br>(0.028) | 0.413<br>(0.020) | 0.444<br>(0.011) | 0.506<br>(0.006) |
| DyScore-RF | 0.845<br>(0.002) | 39.3<br>(1.2) | 35.3<br>(0.6) | 28.2<br>(0.4) | 19.3<br>(0.8) | 10.3<br>(0.3) | 6.2<br>(0.1) | 0.559<br>(0.004) | 0.486<br>(0.006) | 0.453<br>(0.009) | 0.478<br>(0.010) | 0.535<br>(0.009) |
| DyScore-DT-XGBoost | 0.848<br>(0.002) | 42.9<br>(1.5) | 37.9<br>(0.4) | 30.1<br>(0.5) | 20.3<br>(1.1) | 10.5<br>(0.2) | 6.3<br>(0.1) | 0.617<br>(0.002) | 0.526<br>(0.008) | 0.480<br>(0.012) | 0.495<br>(0.012) | 0.546<br>(0.011) |

#### Evaluate on the DUD-E test set with target-unaware split

This experimental setting was designed to represent the case where the targets for scoring do not have known active ligands, and the targets themselves are also novel to the scoring function. DyScore was trained and validated based on the 90% of targets in the DUD-E data set, and the rest 10% of targets (11 in total) were used for testing. With no shared targets between training (or validation) and test sets, this experimental setting was suitable to evaluate the scoring function performance for the newly discovered target with little knowledge about its possible active ligands. The comparison results of our DyScore and the other methods were summarized in **Table 2.** Our method achieves AUC of 0.755, EF-top3 of 20.7 and BEDROC (321.9) of 0.255, comfortably surpassing the others again. Closer inspection of the table shows that the prediction performances of a scoring method over three random test sets differ a lot, which are indicated by the larger standard deviations compared with results in **Table 1,** especially for the early recognition metrics. This diversity might be related to the fact that the model performances vary from target to target, and there exist broad differences in the prediction difficulty between easy and challenging targets. The performance is partially affected by how many challenging or easy targets are randomly split into the test set. Another possible reason is that target-unaware split likely introduces large data distribution differences between training and test sets, which makes some testing targets out of training distribution challenging to be predicted accurately. Thus, increasing and diversifying the training data are promising to improve the DyScore performance further. **Figure 5** illustrates the distribution and median of AUC, EF-1%, BEDROC (321.9), and EF-top3 for different methods, where each gray dot represents a target in the test set. The comparison results on the validation set can be found in **Supplementary Table S2** and **Figure S5**.

**Table 2:** Comparison of DyScore with SOTA scoring functions on the DUD-E test set generated by the target-unaware split. All the methods are evaluated on the three independent random split test sets and the mean value is reported as well as the standard deviation in parentheses.

| Methods | AUC | Enrichment Factor |  |  |  |  |  | BEDROC |  |  |  |  |
| --- | --- | --- | --- | --- | --- | --- | --- | --- | --- | --- | --- | --- |
|  |  | Top 3 | Top 5 | 1% | 2% | 5% | 10% | 321.9 | 160.9 | 80.5 | 32.2 | 16.1 |
| **DyScore** | **0.755**<br>**(0.024)** | **20.7**<br>**(3.2)** | **22.4**<br>**(4.7)** | **12.1**<br>**(1.2)** | **9.2**<br>**(0.8)** | **5.8**<br>**(0.7)** | **4.1**<br>**(0.5)** | **0.255**<br>**(0.020)** | **0.223**<br>**(0.014)** | **0.216**<br>**(0.015)** | **0.256**<br>**(0.024)** | **0.321**<br>**(0.031)** |
| Vina | 0.706<br>(0.027) | 16.8<br>(5.9) | 15.9<br>(7.1) | 10.0<br>(1.6) | 7.6<br>(1.6) | 5.1<br>(1.1) | 3.5<br>(0.6) | 0.211<br>(0.014) | 0.184<br>(0.023) | 0.182<br>(0.031) | 0.218<br>(0.043) | 0.275<br>(0.051) |
| DSX | 0.662<br>(0.012) | 10.7<br>(6.4) | 10.5<br>(3.5) | 5.6<br>(1.4) | 4.9<br>(0.9) | 3.6<br>(0.4) | 2.7<br>(0.3) | 0.110<br>(0.016) | 0.103<br>(0.019) | 0.110<br>(0.019) | 0.148<br>(0.018) | 0.202<br>(0.019) |
| RF-Score-v3 | 0.662<br>(0.037) | 7.7<br>(2.8) | 8.5<br>(2.2) | 4.0<br>(1.6) | 3.7<br>(1.4) | 3.0<br>(0.6) | 2.4<br>(0.4) | 0.075<br>(0.033) | 0.074<br>(0.030) | 0.083<br>(0.030) | 0.121<br>(0.033) | 0.177<br>(0.036) |
| X-Score | 0.658<br>(0.020) | 9.2<br>(3.3) | 7.0<br>(2.6) | 4.5<br>(2.1) | 3.8<br>(1.4) | 2.9<br>(0.7) | 2.3<br>(0.3) | 0.079<br>(0.032) | 0.077<br>(0.030) | 0.085<br>(0.030) | 0.120<br>(0.033) | 0.172<br>(0.033) |
| NNScore-v2 | 0.591<br>(0.018) | 1.3<br>(1.1) | 1.9<br>(1.8) | 2.5<br>(0.9) | 2.4<br>(0.6) | 2.0<br>(0.4) | 1.8<br>(0.3) | 0.043<br>(0.014) | 0.044<br>(0.013) | 0.052<br>(0.014) | 0.081<br>(0.018) | 0.128<br>(0.020) |
| Score2 | 0.608<br>(0.005) | 2.4<br>(0.9) | 2.2<br>(0.3) | 2.4<br>(0.4) | 2.1<br>(0.3) | 2.1<br>(0.2) | 1.9<br>(0.1) | 0.038<br>(0.007) | 0.041<br>(0.006) | 0.050<br>(0.006) | 0.082<br>(0.007) | 0.132<br>(0.008) |

**Fig. 5:**
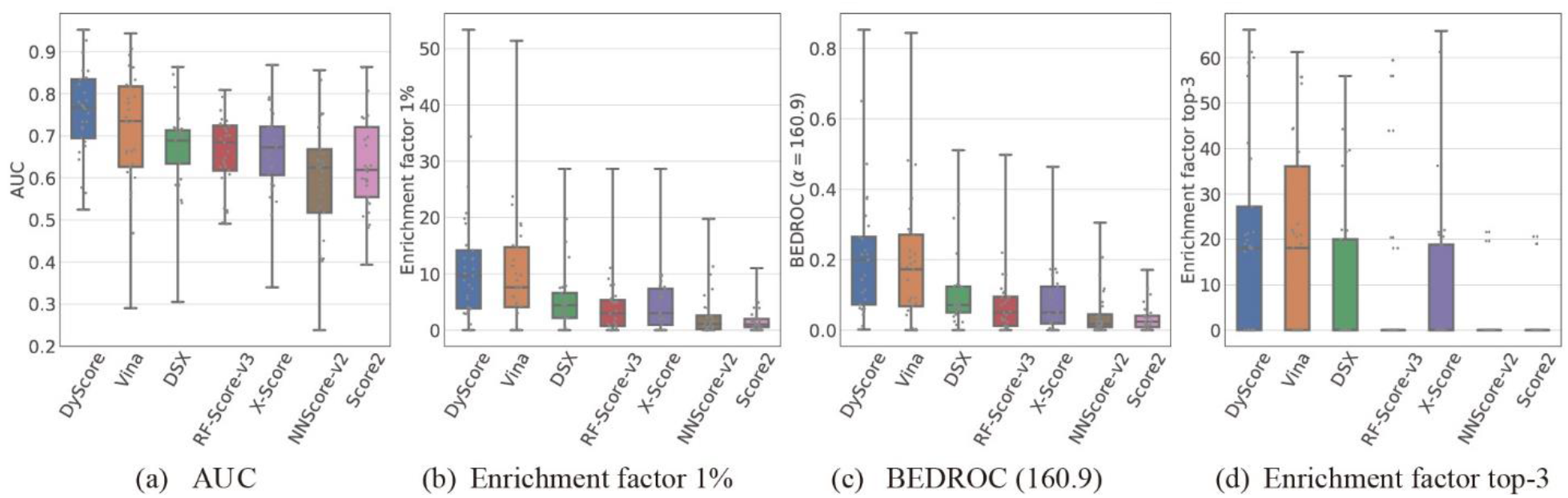
Comparison of different scoring methods on the DUD-E test set generated by target-unaware split. Each gray dot represents a target.

#### Evaluate on the DEKOIS 2.0 external test set

We further evaluated our DyScore model using the external benchmark test set DEKOIS 2.0^23^. As described in Section 2.1, we removed four targets (A2A, HDAC2, PARP-1, and PPARA) from the DEKOIS 2.0 test set also present in the DUD-E data set DyScore had been trained on to prevent any potential data leakage. Since the DEKOIS 2.0 and DUD-E datasets were constructed using different strategies for collecting active compounds and generating decoy compounds and have different active/decoy ratios, we evaluated the performance of our DyScore model by using other methods as references. **Table 3** shows the comparison results of our proposed DyScore and the six SOTA methods on the unseen 77 DEKOIS targets. We found that DyScore remarkably outperforms the competitors over all measurements. On the most critical early recognition metrics EF and BEDROC, our DyScore achieves EF-top3 of 10.5 and BEDROC (321.9) of 0.285. Arguably the most commonly used docking software, AutoDock Vina could be considered a recognized reference point for estimating both the performance and generalization ability of a model. When compared to Vina, the second-best method, our DyScore obtains the relative improvements of 69.4% and 52.4% on these two metrics, respectively. The increase of relative improvements on DEKOIS 2.0 test set over the target-unaware split DUD-E dataset (with relative improvement 23.2% and 20.9%) suggests that DyScore shows a more distinct improvement recognition capacity in a dataset that is tougher to predict. The comparison results adequately demonstrate that the enhanced performance originated from the two proposed features in DyScore is robust and can be well generalized to the external data set. In addition to comparing methods on their averaged performances over the targets, we also used the box and whisker plot to summarize and illustrate the detailed distribution of AUC, EF-1%, BEDROC (321.9), and EF-top3 in **Figure 6**. Closer inspection of the EF-top3 plot shows that the median value of DyScore is around 10, but the others are all close to 0. This indicates these competing approaches are unable to find any true binders among their top3 ranked predictions for half of the targets in the DEKOIS 2.0 dataset, presenting a significant challenge in the experimental drug discovery process, especially in *de novo* design. DyScore predictions, however, have a much higher accuracy and show great potential for use in computational drug development protocols. The 77 per-target ROC curves generated by DyScore on the DEKOIS 2.0 dataset are shown in **Supplementary Figure S8**.

**Table 3:** Comparison of DyScore and other SOTA methods on the DEKOIS 2.0 data set.

| Methods | AUC | Enrichment Factor |  |  |  |  |  | BEDROC |  |  |  |  |
| --- | --- | --- | --- | --- | --- | --- | --- | --- | --- | --- | --- | --- |
|  |  | Top 3 | Top 5 | 1% | 2% | 5% | 10% | 321.9 | 160.9 | 80.5 | 32.2 | 16.1 |
| **DyScore** | **0.702** | **10.5** | **9.6** | **7.5** | **6.2** | **4.3** | **3.3** | **0.285** | **0.244** | **0.216** | **0.227** | **0.277** |
| Vina | 0.631 | 6.2 | 6.4 | 4.8 | 4.0 | 2.8 | 2.3 | 0.187 | 0.159 | 0.140 | 0.151 | 0.192 |
| DSX | 0.590 | 4.2 | 4.3 | 3.4 | 2.7 | 2.3 | 1.9 | 0.122 | 0.108 | 0.100 | 0.116 | 0.157 |
| RF-Score-v3 | 0.594 | 3.0 | 3.3 | 3.1 | 2.7 | 2.4 | 2.0 | 0.102 | 0.096 | 0.095 | 0.118 | 0.164 |
| X-Score | 0.597 | 3.6 | 3.7 | 3.5 | 3.2 | 2.5 | 2.0 | 0.115 | 0.109 | 0.107 | 0.125 | 0.166 |
| NNScore-v2 | 0.567 | 1.6 | 2.1 | 2.0 | 1.8 | 1.6 | 1.4 | 0.059 | 0.060 | 0.061 | 0.078 | 0.114 |
| Score2 | 0.595 | 1.3 | 2.0 | 2.2 | 2.3 | 2.2 | 2.0 | 0.061 | 0.067 | 0.075 | 0.104 | 0.152 |

**Fig. 6:**
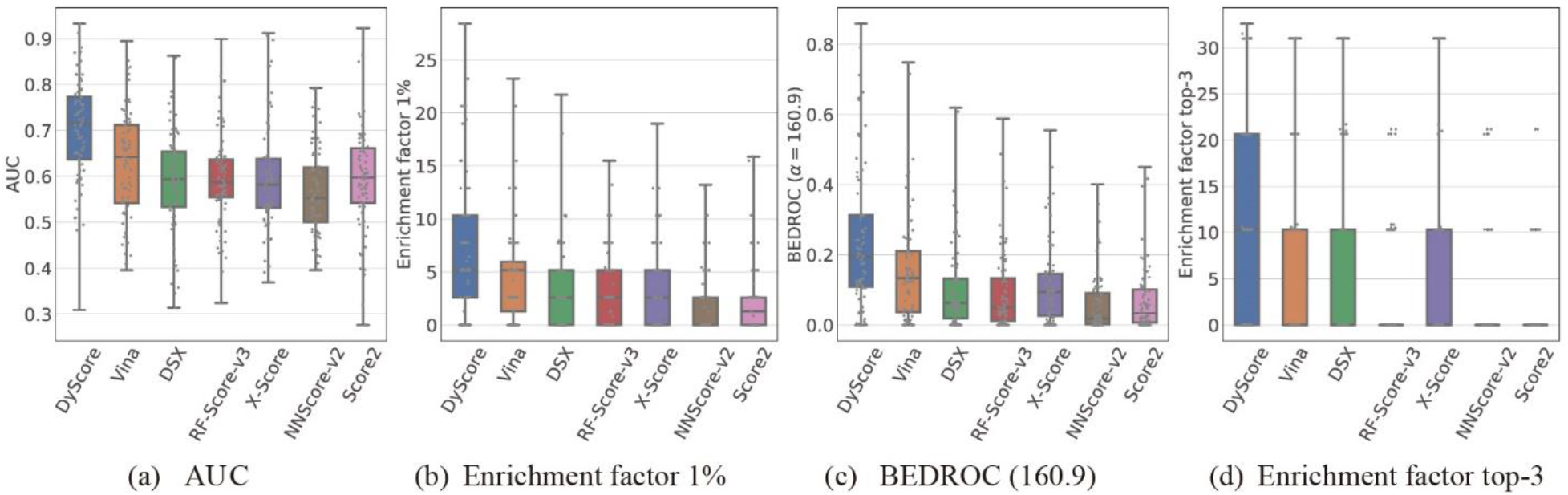
Comparison of different scoring methods on the DEKOIS 2.0 data set. Each gray dot represents a target from DEKOIS dataset.

We also analyzed DyScore’s prediction distribution on the DUD-E test set generated by target-aware split and DEKOIS 2.0 data set. The probability density function was estimated using a Gaussian kernel, and the predictions fall in the range [0, 1], representing the predicted confidence score of a true binder. As shown in **Figure 7,** for both data sets, prediction values for decoy samples are mostly concentrated in a small range around 0, which is as expected. Although some predictions for active samples are relatively small, in the high confidence range, which is amplified inside the figure, density for active predictions is much higher than the decoy predictions. This indicates that selecting samples from our top ranked predictions, we have many more chances to get a true binder, which is as same as early recognition.

**Fig. 7:**
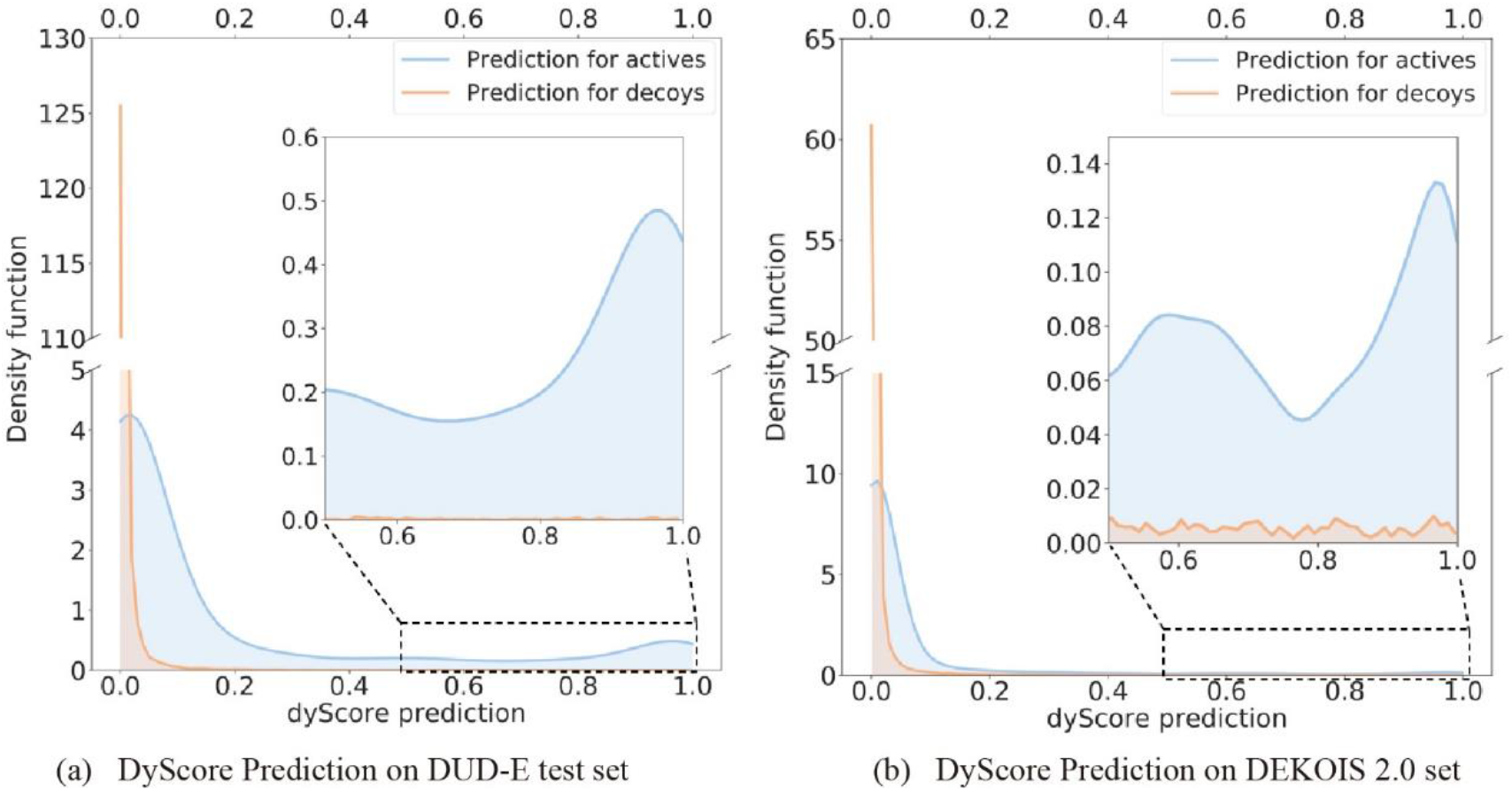
The estimated density functions of DyScore prediction on DUD-E test set with target-aware split (left) and DEKOIS 2.0 benchmark set (right)

### 3.2 The Proposed Dynamic Properties Significantly Improve Performance of the Model

Classical scoring functions can be roughly categorized into three main classes: force field-based, knowledge-based, and empirical^46^. Scoring functions are not accurate due to their substantial rough approximation or even omittance of many aspects of protein-ligand binding. For instance, the intrinsic dynamic of the protein-ligand binding state is usually disregarded in scoring functions because these prediction approaches are limited to obtaining information from the static structure of a complex. Despite that, it would be unrealistic to estimate the dynamic property of protein-ligand binding by sampling the possible binding complexes in different dynamic states with a rapid scoring function. However, although the static snapshot of complex structure does not explicitly present the mobility of ligand in the binding site, the potential fluctuation of ligand should be relevant to its interactions with protein residues. So, it would be possible to estimate the dynamic properties of protein-ligand binding based on the static snapshot of a complex structure. The importance of the two proposed binding features, the stability score and matching score, was measured by performing the permutation importance experiment. Specifically, a test set was permuted by replacing the evaluated feature with random noise, i.e., randomly shuffle its values across different samples^47^. Then we used the DyScore model, which was pre-trained on the original 14 features, to predict this permutated test set, and we measured the decrements of the model’s performance. Consistent with the experimental settings described in Section 3.1, the permutation importance experiments were still conducted on the three different data sources, including two test sets respectively generated by target-aware and target-unaware data splits and DEKOIS 2.0 external test set. For each data set, five randomly permutated sets were generated individually for stability score, matching score, and both of them to reduce the bias.

As **Figure 8** shows, across all metrics, including AUC, EF, and BEDROC, the permutation of both lock-key matching score and protein-ligand stability score, named as double permutation, significantly downgrades the performance of the model, and this comparison result keeps consistent across all the different test sets. In **Figure 8a**, by comparing the double permutation with the non-permutation results, we can find that the two proposed features jointly improve the AUC score from 0.792 to 0.858, and bring about remarkable relative improvements on the early recognition metrics, where 86.1% for EF-1%, 79.5% for BEDROC (160.9) and 79.7% for EF-top3. This implies that the superior performance of DyScore is highly dependent on the two new features. Additional, double permutation on the test set induced the decrements of EF-1% by 14.9, which is approximating to the sum of effects for individual permutation of stability score and matching score (7.8 + 4.8 = 12.6). Considering that the BEDROC (160.9) on the stability score permutation set and matching score permutation set shows a similar additive effect, it could be speculated that the stability score and matching score are likely to be orthogonal to each other to a large extent. It is intuitive to expect that the two features may partially overlap because both features are designed to describe the dynamic properties of protein-ligand binding. However, the matching score is a measurement of the local gaps between protein and ligand, which is mainly related to the restriction of local mobility for ligand inside the binding site. The stability score, however, is mainly related to the resistance of the external perturbation caused by the Brownian motion of solvent molecules, so the matching score and stability score are expected to contribute to the model’s performance independently to a large extent. It can also be seen from the figure that the permutation on stability score consistently geunerates better results than permutation on the matching score, which indicates that the matching score contributes more to the DyScore’s performance. **Figure 8b** and **Figure 8c** illustrate the permutation importance results on DUD-E test set with target-unaware split and DEKOIS 2.0 data set. Similar feature importance trends can be observed from them, which empirically verifies the significant contribution of our two proposed dynamic properties to the DyScore performance.

**Fig. 8:**
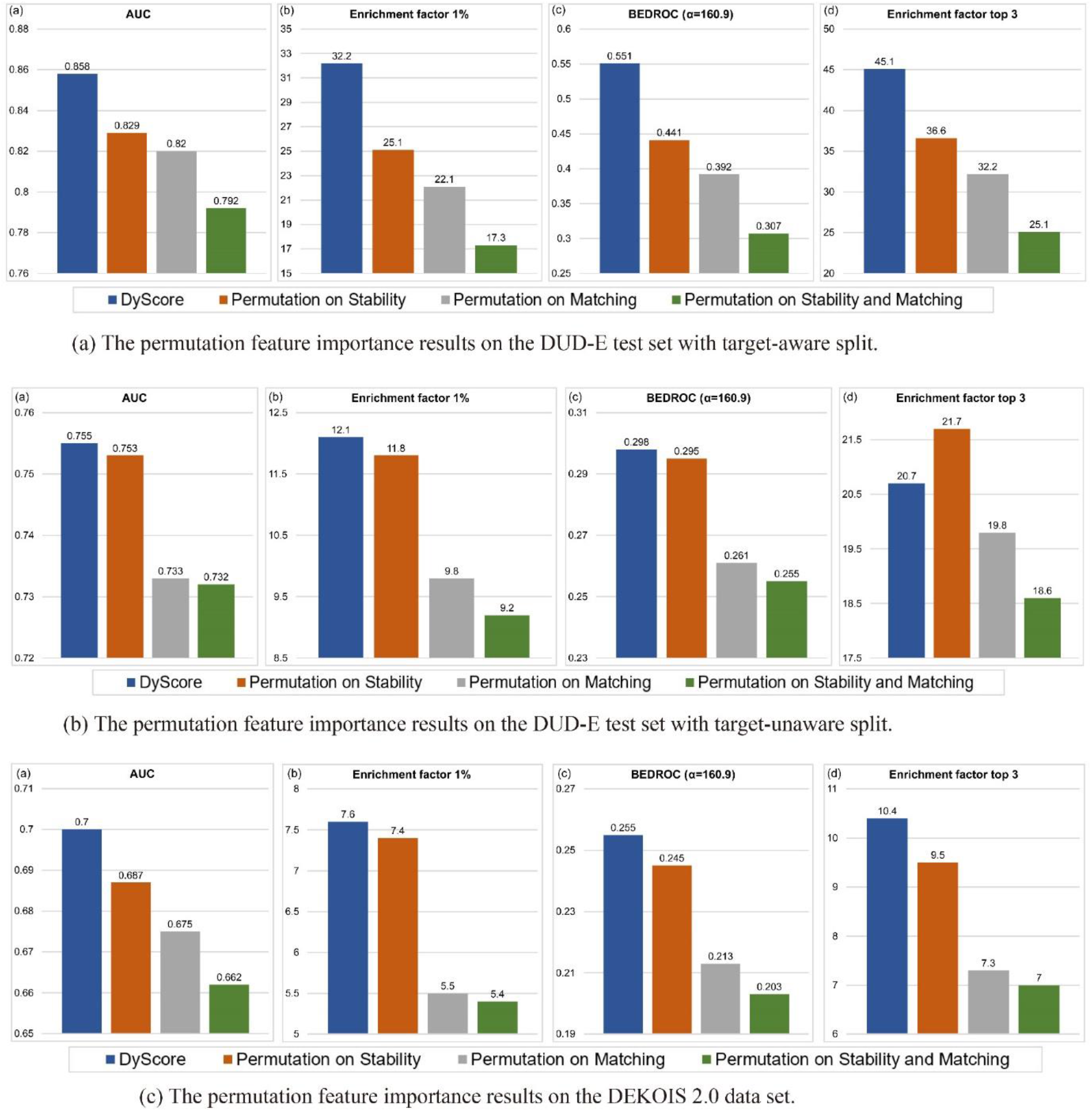
The permutation feature importance results of the stability and matching scores.

### 3.3 Difference in Feature Selection for Molecular Similarity-based and Interaction-based Learning Model

It is well-known that compounds with similar topological structures usually suggest their potential similar bioactivity to particular protein target^48^. Furthermore, it is also generally recognized that known active compounds for a particular protein target usually could be clustered into one or a few groups, which implies that active compounds of each target are usually limited in a tiny region of chemical space on a target-by-target basis. Especially, according to the DUD-E dataset construction strategy, the structural similarity between decoys and actives are minimized. Thus, active and decoy ligands of the same target protein are expected to show distinct characters in topological structural clustering based on their fingerprints. **Figure 9** is a t-SNE map depicted the 2-D feature space of the ligand fingerprints of 3 targets from DUD-E dataset, which demonstrate the potential basis of incorporating ligand information in the training set. The dimension of fingerprints is reduced from original 1024 to 50 by principal component analysis (PCA) method and sequentially reduced to 2 by the t-SNE algorithm. Dots representing decoy ligands are widely dispersed in the feature space; on the contrary, the active ligands are highly clustered. It implies that if a machine learning model explicitly or implicitly uses ligand structure information, e.g., fingerprint, as its training inputs, it likely only leverages this obvious difference between ligands to distinguish actives and decoys and easily obtains a high prediction accuracy when testing on some similar ligands. Recently, several machine learning and deep learning-based approaches have been proposed for compound pose classification (binder or non-binder) and achieved better performances on the DUD-E dataset, compared with the conventional methods, such as empirical or knowledge-based scoring functions^49–51^. Such similarity-based machine learning methods are expected to show a higher success rate in finding binders that are similar to known binders in practical drug discovery projects. However, the weak generalization abilities of certain machine learning models trained on the DUD-E dataset have already been reported by several groups^52, 53^, where these methods perform comparable or worse than the Vina score if the test ligands share less similar topological structures with the training samples. On the contrary, a machine learning method that only considering the protein-ligand interaction information, namely the interaction-based method, is expected to learn more basic physical-chemistry rules involved in protein-ligand binding, which could have better generalization abilities in predicting novel binders that dissimilar with known binders. As a result, we will not compare the similarity-based and interaction-based learning models on one page because they fit most for different application scenarios.

**Fig. 9:**
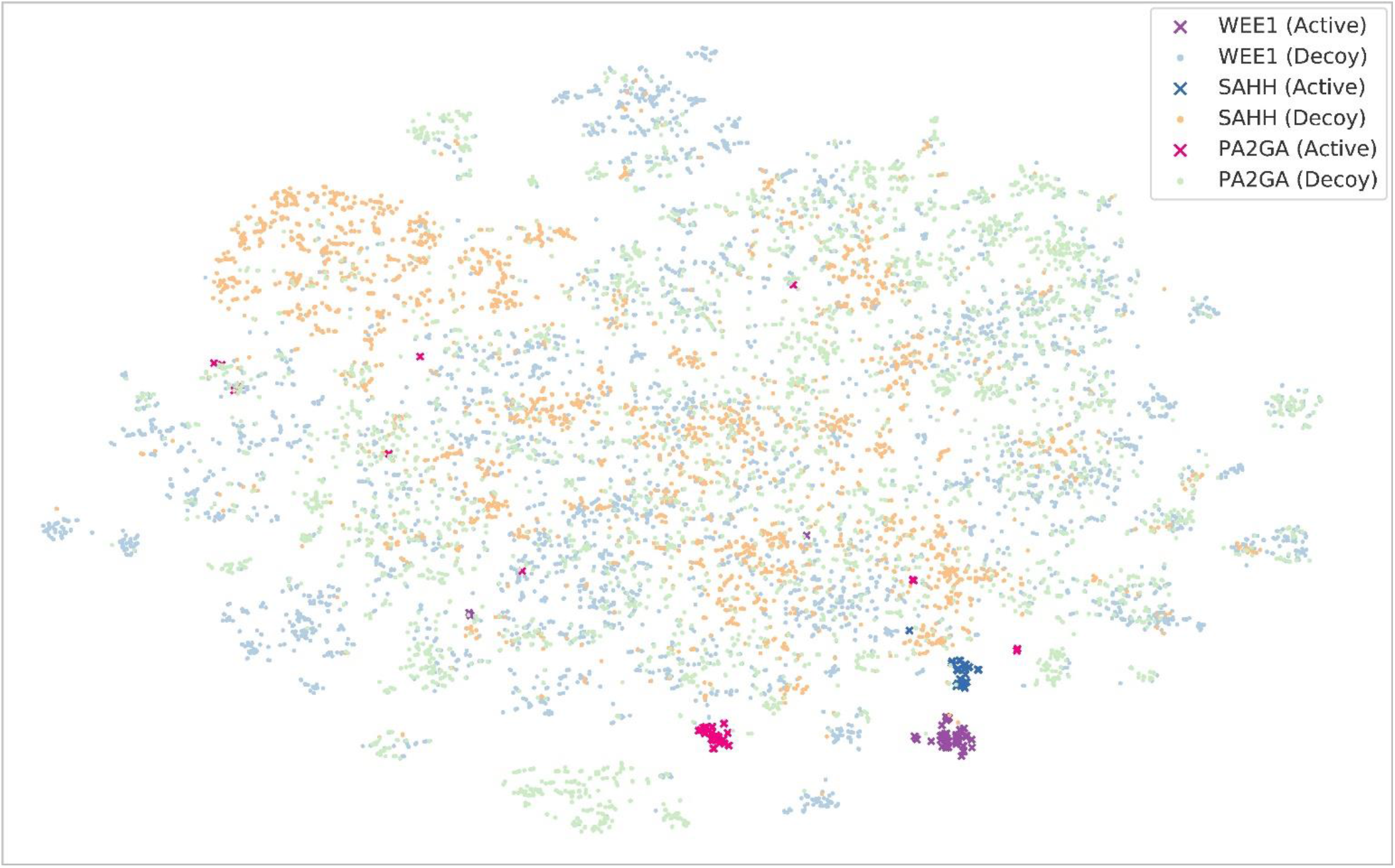
Ligand fingerprint visualization for three targets (i.e, WEE1, SAHH and PA2GA) in two-dimensional space. Starting from the original 1024-dimensional ligand fingerprint, we first performed PCA to reduce the dimension to 50, then ran the t-SNE algorithm to further reduce to two dimensions. The cross and dot respectively represent the actives and decoys.

DyScore is developed based on the concept of the interaction-based model, which is optimized to achieve better generalization ability. To avoid inducing information from structural similarity, we only use the features that can characterize protein-ligand interaction instead of the ligand property to train the DyScore model in this study. In short, any information that only depends on the ligand itself should not be used as training input. Note that the DUD-E dataset was constructed based on the resemble of distribution for four physical properties (including molecular weight, logP, number of rotatable bonds, and hydrogen bond donors and acceptors), it would be safe to use only one of the four physical properties in training data because the active and decoy are statistically indistinguishable if only one property alone is investigated. In DyScore, the number of rotatable bonds is selected as one of the input features because it is highly related to conformational entropy loss of ligand during the binding process and tends to be the most binding relevant property among the four physical properties. The remaining input features are composed of several classical scoring functions not trained on the DUD-E dataset to avoid any potential bias.

Although we do not emphasize the similarity-based method in this work because the performance for published similarity-based methods is already close to the ceiling, we still developed a modified similarity-based DyScore model by integrating the molecular fingerprints (MF), named DyScore-MF. Molecular fingerprints are widely used in encoding the structure information of molecules. In addition to the 14 binding features mentioned in Sections 2.2 and 2.3, we utilized path-based FP2 fingerprints with 1024 binary digits (bits) generated by OpenBabel to encode the fragment and local topology information in the molecule^41^. Based on the combined 1038 features, we trained a new XGboost model with the same parameter settings described in Section 2.5 on the DUD-E dataset with the target-aware splitting and compared it with the original DyScore model. According to **Figure 10**, we can find that after integrating the MF, DyScore-MF can easily achieve nearly 1.0 AUC score, and its superior performances can also be observed on the other metrics. The active and decoy ligands of a specific target protein tend to show distinct characters in their fingerprints, and the active compounds usually could be clustered to a few structural pattens. Therefore, introducing MF to a machine learning model could significantly improve its accuracy in identifying active compounds that are similar to any active compound in the training dataset. This implies that the DyScore-MF is feasible to identify nearly all active binders from the DUD-E dataset, which demonstrates the superiority of the similarity-based method in finding binders that similar to known compounds in the training data set. Unsurprisingly, the integrated MF features can enhance the DyScore-MF’s performance on the DUD-E dataset. Although the input of DyScore-MF model also contains 14 protein-ligand interaction-based features used in DyScore, the DyScore-MF model would be primarily relying on the MF part considering its significant performance improvement comparing with DyScore. We believe similarity-based methods like DyScore-MF could be very effective in identifying new binders based on known binders; however, the interaction-based method is expected to be more helpful in identifying diverse novel binders based on learned physical-chemistry rules of protein-ligand interaction. To demonstrate our method can effectively capture the protein-ligand interaction instead of relying on the ligand similarity, we solely consider DyScore as our proposed benchmark method instead of DyScore-MF. For a fair evaluation, we only compare DyScore with other interaction-based methods. All the baseline approaches compared in the main text are carefully selected to make sure no competitor is built based on the ligand structural similarity.

**Fig. 10:**
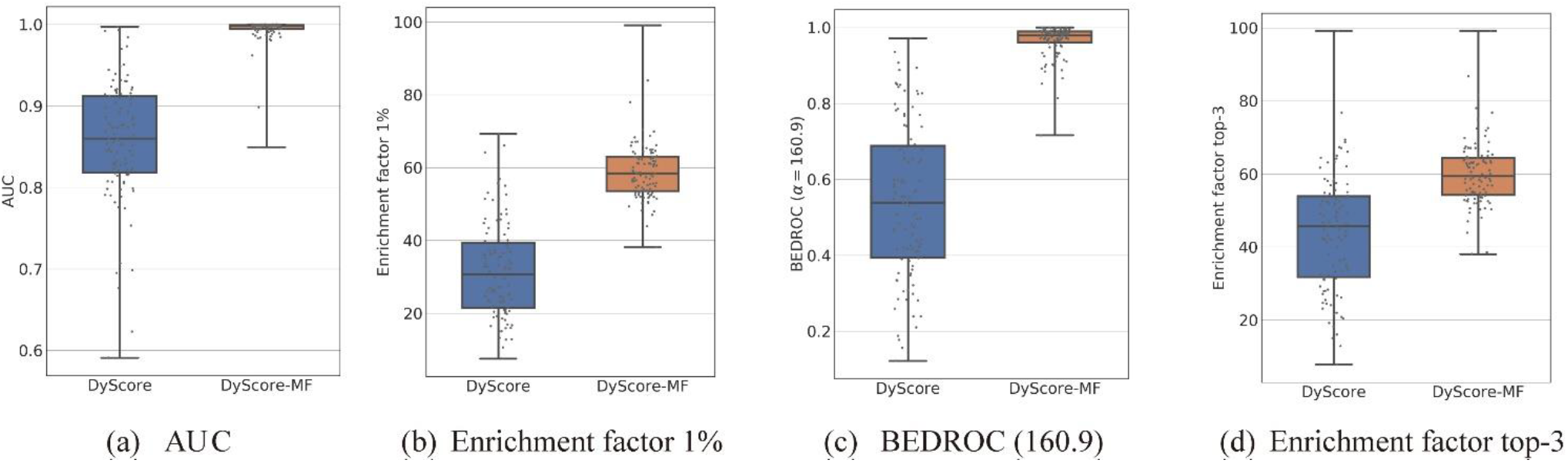
Comparison of interaction-based DyScore and similarity-based DyScore-MF methods on the DUD-E test set generated by target-aware split.

To further demonstrate the validity of DyScore, we have compared the performance of DyScore and DyScore-MF on the second external test set LIT-PCBA with total twelve SOTA methods. To illustrate the performance of similarity-based DyScore-MF, we have added six recently published SOTA methods with removing the exclusion of similarity-based model. Due to the relatively weak binding affinity of active compounds from the HTS based LIT-PCBA dataset, DyScore, DyScore-MF and all SOTA methods shows significant performance degradation comparing with they work in DUD-E dataset. However, although the EF1% of interaction-based DyScore model only surpass 10 in 12 SOTA methods, EF1% of the similarity-based DyScore-MF still outperform all other SOTA methods including many similarity-based approaches, which suggests that the DyScore-MF also benefits from other features proposed in DyScore model.(Supplementary Table S4) The interaction-based DyScore model is expected to have better generalization ability than similarity-based model, especially for some novel targets lacking prior knowledge for ligand. However, similarity-based model may have better performance for common targets with plenty of prior knowledge for ligand. Therefore, both DyScore-MF and DyScore model would be useful in virtual screening practice.

### 3.4 Comparison of DyScore and Other Methods Learning from the Same Features

This section evaluates different machine learning and deep learning methods trained on the same 14 binding features. Compared with all the competing algorithms shown in the second part of **Table 1,** DyScore with RF-based XGBoost algorithm achieves the best performance on all the evaluation metrics, especially on the early recognition metrics EF and BEDROC. The second-best model, DyScore-DT-XGBoost, uses the same boosting strategy as Dyscore but uses decision trees as base learners instead. The major difference is that an RF model consists of multiple single decision trees independently trained with bootstrap samples. Additionally, to further increase the trees’ diversity in a RF model, a random subset of features is considered when splitting at each tree node. The ensemble design and introduced randomness make the RF model more potent and robust than a single decision tree. As we expected, DyScore respectively achieves the 5.1% and 4.4% relative improvements on the EF-top3 and BEDROC (321.9) metrics, compared with DyScore-DT-XGBoost. Another competing model, DyScore-RF, is built purely based on the random forest model. Compared with it, our DyScore model sequentially uses the boosting training policy to add additional new RF models to complement the previously built ones, and the original bagging and random feature subsampling strategy of RF are still incorporated. Empirically, DyScore respectively enhances the EF-top3 and BEDROC (321.9) metrics from 39.3 to 45.1 and 0.617 to 0.644. Comparing with DyScore-MLP, DyScore largely outperforms it by relative improvements of 28.1 and 27.3 on the EF-top3 and BEDROC (321.9).

### 3.5 Early Recognition in De Novo Design

As researchers usually use a VS approach to select tens to hundreds of compounds for the experimental test, the most meaningful EF and BEDROC focus on the top 0.5% or 1% of compounds in a typical VS study. However, the EF of a few top ranked prediction are more important for *de novo* drug design, because very few compounds are considered for exploration in *de novo* drug design due to the high cost of synthesizing compounds. As a result, the accuracy of top batch compounds predicted by the model is crucial in *de novo* design. According to **Figure 11**, although the performance of DyScore and six SOTA methods are improved to some extent when focusing on top batch predictions, DyScore shows clearly superior performance in identifying active compounds from top batch predictions. Also noteworthy, DyScore demonstrates a more robust performance across 102 protein targets; on the contrary, EF-top3 of other methods fall to zero for nearly half of the protein targets, a sign of weak generalization ability. Furthermore, the EF-top3 for DyScore is 45.1, specifically, average 224.3 active compounds were identified in all 306 predicted candidates (top3 for 102 targets), which implies a superior 73.3% success rate for top ranked compounds. **Figure 12** and **Figure 13** respectively show the similar trend on DUE-E test set with target-unaware split and DEKOIS 2.0 external dataset. More results on the DUD-E validation sets are shown in **Supplemental Figure S4** and **S6**. The remarkable improvement of DyScore on the performance of EF - top3 implies its promising application in *de novo* design.

**Fig. 11:**
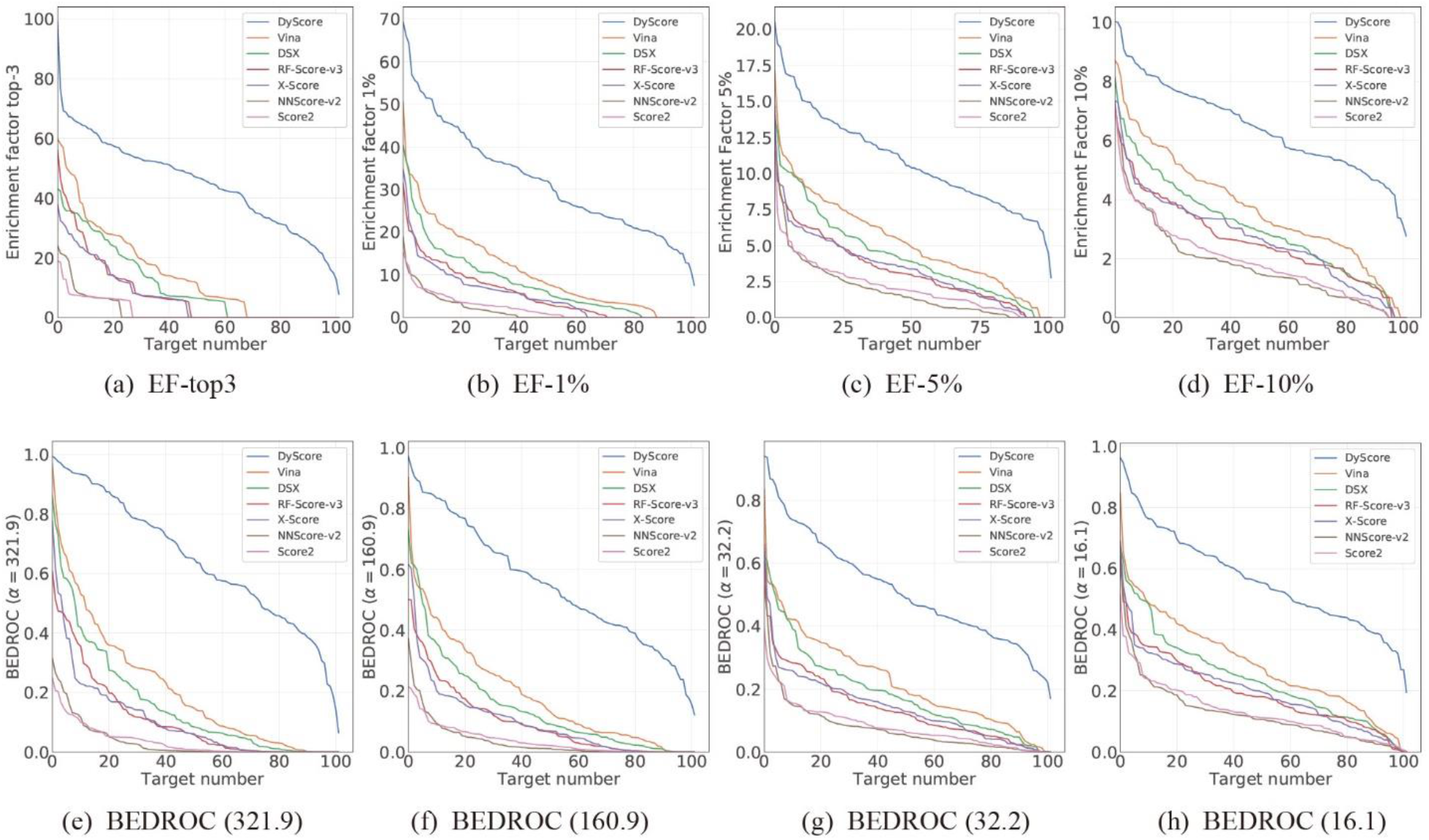
Enrichment factor and BEDROC to evaluate the performance of different scoring methods with respect to the target number. The metrics are measured on the DUD-E test set generated by the target-aware split.

**Fig. 12:**
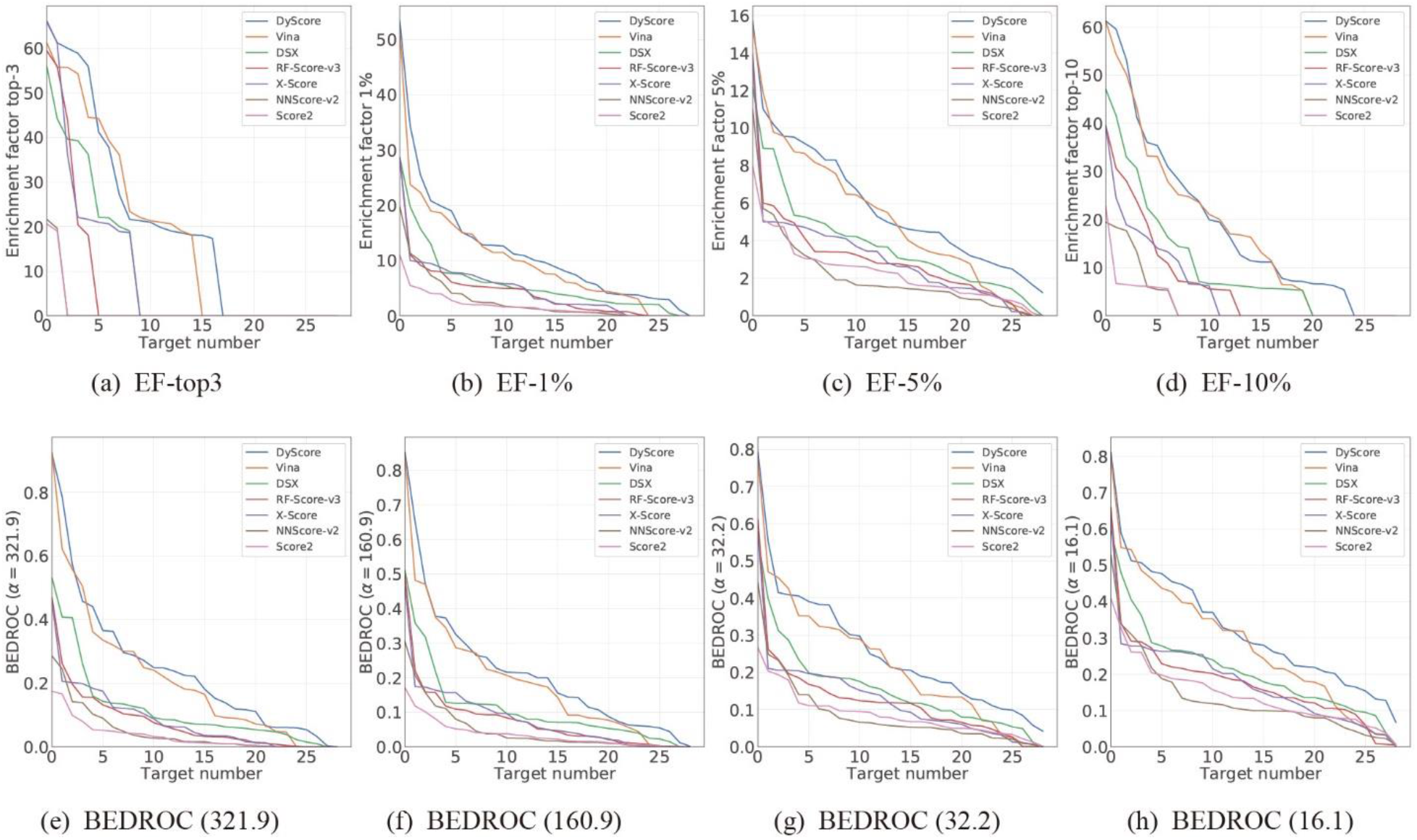
Enrichment factor and BEDROC to evaluate the performance of different scoring methods with respect to the target number. The metrics are measured on the DUD-E test set generated by the target-unaware split.

**Fig. 13:**
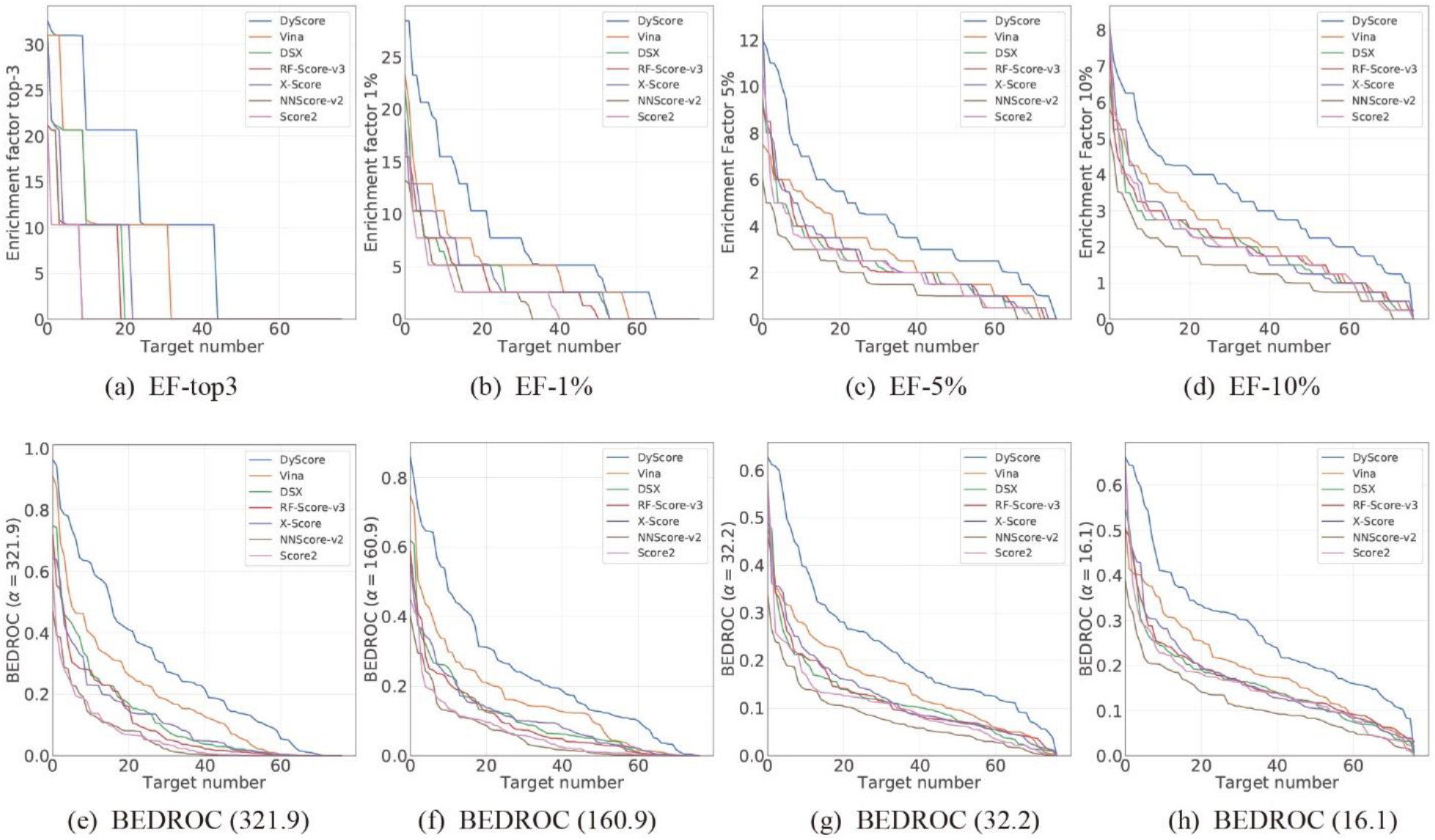
Enrichment factor and BEDROC to evaluate the performance of different scoring methods on DEKOIS 2.0 data set with respect to the target number.

### 3.6 Computational Time of DyScore

Total 14 input features were needed for the interaction based DyScore model. For similarity-based DyScore-MF model, an additional FP2 binary bit vector is also needed. The computational time for generating the input features were listed in Supplementary Table S5. The main computational cost of DyScore is in the step of running different scoring functions, so the computational cost is in the approximate order of magnitude to the slowest scoring function used in the model. Besides, the calculation of two dynamic features takes 18% computational time of the whole process. In conclusion, the computational cost of DyScore is still comparable with the typical scoring functions.

## 4 Conclusion

In this study, we proposed two novel features for estimating the dynamic properties of the protein-ligand binding process based on the static structure of the protein-ligand complex. The geometry matching score was designed to measure the local gaps between protein and ligand, which is associated with the restriction of local mobility for ligand inside the binding site. The dynamic stability score was proposed to estimate the stability of a ligand binding to a protein, which is related to the resistance of the perturbation caused by the thermal fluctuation and random hitting from solvent molecules. Utilizing these two features and several classical scoring functions, we developed the DyScore binary classification model that uses the XGBoost algorithm to classify compound poses as binders or non-binders. DyScore was trained and evaluated on the widely used benchmark DUD-E dataset and externally tested on both DEKOIS 2.0 and LIT-PCBA dataset. Extensive experimental results demonstrated that the DyScore model surpasses current SOTA scoring methods by a large margin and the inclusion of both our proposed novel features significantly improved the ranking power and performance of the DyScore model. Besides, the DyScore-derived similarity-based DyScore-MF model also surpasses SOTA methods that trained with ligand similarity information, which could be used for screening targets with sufficient prior knowledge for ligand. More importantly, the DyScore model showed superior performance in early recognition with an average of 73.3% of the top three ranked compounds for each protein target proving active. This is promising for improving the success rate of virtual screening and *de novo* drug design.

## 5 Data and Software Availability

The standalone version of DyScore and DyScore-MF are freely available to all at the following website: https://github.com/YanjunLi-CS/dyscore. The Docker container image of DyScore could be built with the pre-prepared Dockerfile, which could automatically install all dependents and set up running environments. With Docker container, no other dependents or software are required for using DyScore. The docker image should be built in Linux system, but the built docker image could be used in any platforms supported by Docker container. In additional, an optional automatic binding site detection procedure with Cavity^54^ as well as an optional automatically molecular docking function with Autodock Vina^27^ were embedded in DyScore, which enables user to use DyScore without defining binding site and prior docking. The input data generated from docked structures for all the three database (DUD-E, DEKIOS, LIT-PCBA), as well as the output data from DyScore were deposited to google cloud driver (link: https://drive.google.com/drive/folders/1gVRrqpbNRd1_GuPjntVWhStcD8JgWEJn; For description of each items in the dataset, please refer to supplementary/experimental_data directory) in https://github.com/YanjunLi-CS/dyscore. The model prediction and training code with examples were also deposited to supplementary/experimental_data directory.

## ASSOCIATED CONTENT

Supporting Information available. Additional experimental results, as well as figures and tables for detailed metrics evaluating performance of DyScore and DyScore-MF.

