## Supplemental pdf for "DyScore: A Boosting Scoring Method with Dynamic Properties for Identifying True Binders and Non-binders in Structure-based Drug Discovery"

### 1 Additional Results

#### 1.1 Evaluate on the LIT-PCBA external test set

We further evaluated our DyScore model using the external benchmark test set LIT-PCBA. LIT-PCBA datasets contain multiple PDB templates for single targets, so all the metrics are calculated from the protein-ligand complex with the highest predicted active probability for each ligand. As described in Section 2.1, we did not remove duplicate targets from LIT-PCBA for comparison of results with other studies that also kept the duplicates. Since the LIT-PCBA and DUD-E datasets were constructed using different strategies for collecting active compounds and generating decoy compounds and having different active/decoy ratios, we evaluated the performance of our DyScore and DyScore-MF models by using other methods as references. Supplementary Figure S9, S10 and Table S3 shows the comparison results of our proposed DyScore and DyScore-MF on the six SOTA methods. The interaction-based DyScore model has better performance on EF1% compared with all other methods. The similarity-based DyScore-MF model trained with additional fingerprint information shows even higher performance than DyScore. Because all recently developed approaches that tested on LIT-PCBA dataset used the information of ligand structure, we have compared the performance of these methods with DyScore-MF for fairness. DyScore-MF shows better EF1% performance compared with other approaches that were tested on LIT-PCBA dataset. The results demonstrate the validity of using DyScore and DyScore-MF in virtual screening projects. Although DyScore shows advantages compared with other methods, the performance of either DyScore or DyScore-MF on LIT-PCBA dataset is lower than their performance on DUD-E dataset. We think this is mainly due to the different definitions of active compounds in the different datasets. For DUD-E dataset, the active compounds were defined with an affinity threshold of lower than 1uM, and the decoy compounds were defined with an affinity threshold of higher than 30uM. For LIT-PCBA dataset, the average pEC50 or pIC50 of all active compounds is 5.11 (about 7.8uM), and only 3.1% (885 in 10032) of active compounds in LIT-PCBA dataset could achieve 1uM potency. That is, 96.9% of active compounds in LIT-PCBA dataset could not reach the lowest activity threshold of active compounds in DUD-E dataset. So the significant difference in the definition of active compounds may impair the performance of using DyScore to predict the LIT-PCBA dataset because DyScore is trained based on DUD-E dataset. In this case, most "active" compounds in LIT-PCBA dataset with affinity worse than 1uM are likely to be predicted to "decoy" by DyScore, because DyScore would be optimized to fit the active threshold at 1uM. This could explain why DyScore only has a small improvement on EF1% compared with the baseline Score2 function. To prevent over-interpretation of the small difference, we could not make further analysis of the contribution of new features in using DyScore against LIT-PCBA dataset. In the contrast, the similarity-based DyScore-MF model shows a more significant improvement of EF1%, which is possibly due to the intrinsic fuzziness of the similarity-based approach. That is, the similarity-based DyScore-MF is more sensitive to the similarity of ligand and less sensitive to the protein-ligand interaction, which could help to improve the performance even if the definition of "active" and "decoy" is shifted. Besides, the higher EF1% of DyScore-MF compared with other SOTA methods may suggest DyScore-MF also benefits from other features proposed in DyScore model. Furthermore, we could wisely combine the DyScore-MF and DyScore in virtual screening. DyScore-MF could help to identify hits relying on prior knowledge, and DyScore is expected to have a better generalization ability to identify novel compounds with less dependent on prior knowledge.

### 1.2 Distribution of each input feature in DUD-E data set

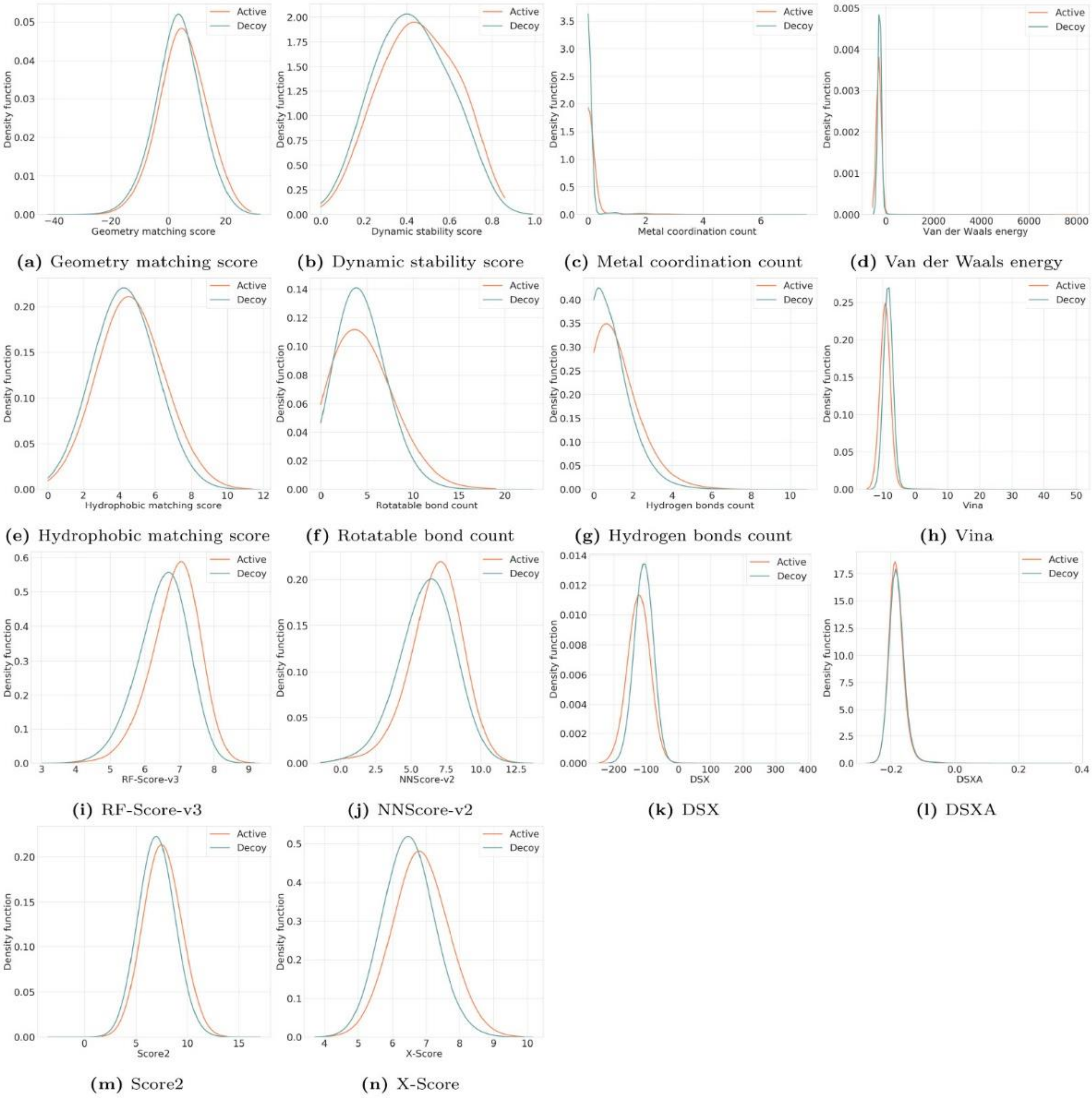


**Figure S1:** Distribution of each input feature in DUD-E data set.

### 1.3 Distribution of each input feature in DEKOIS 2.0 data set

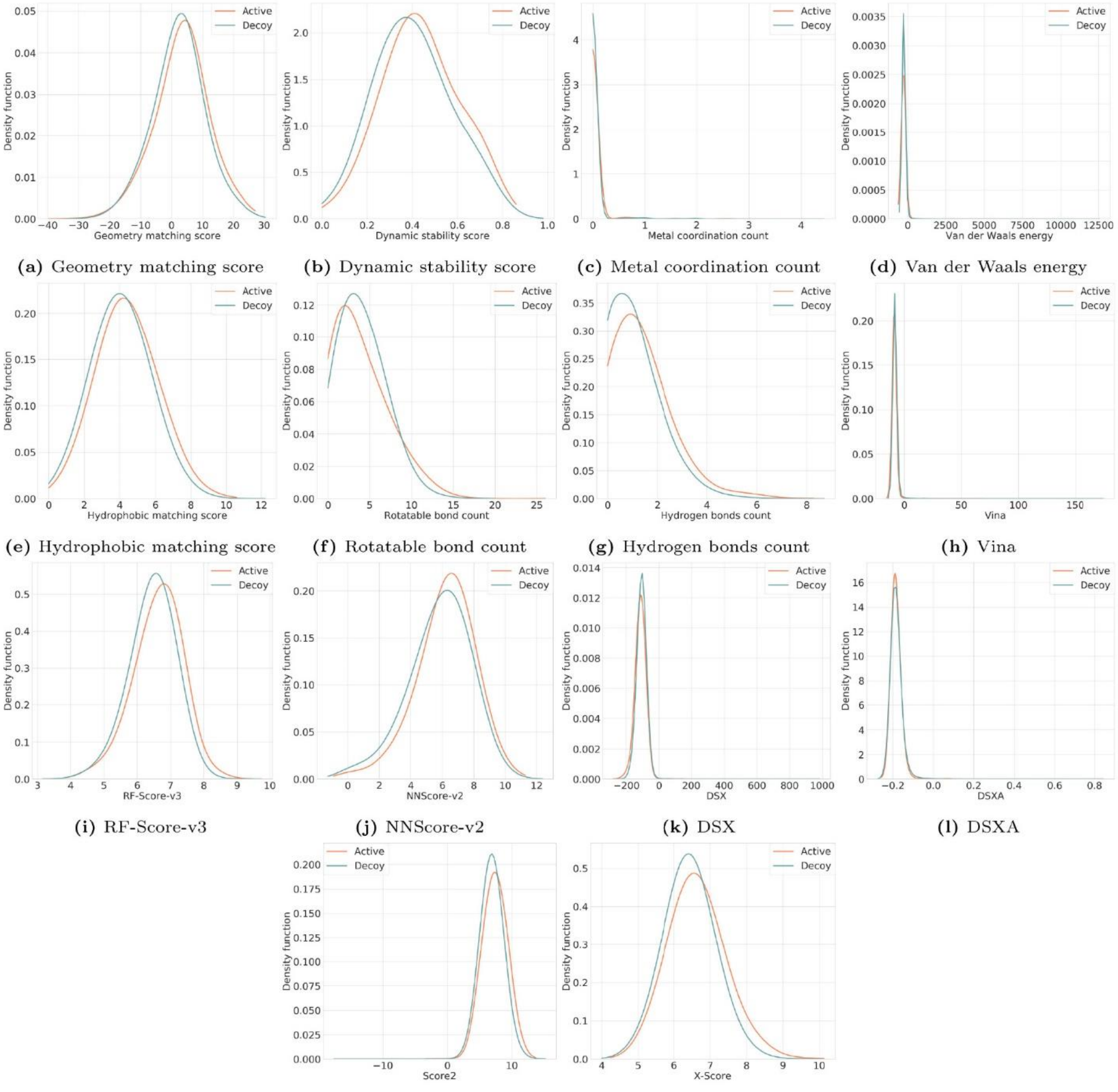


(a) Geometry matching score (b) Dynamic stability score (c) Metal coordination count (d) Van der Waals energy
(e) Hydrophobic matching score (f) Rotatable bond count (g) Hydrogen bonds count (h) Vina
(i) RF-Score-v3 (j) NNScore-v2 (k) DSX (l) DSXA
(m) Score2 (n) X-Score

**Figure S2:** Distribution of each input feature in DEKOIS 2.0 data set

### 1.3 Comparison on the DUD-E Validation Set with Target-aware Split

**Table S1:** Comparison of DyScore with SOTA scoring functions and other methods learning from the same features on the validation set.

| Methods | AUC | Enrichment Factor | | | | | | BEDROC | | | | |
|---|---|---|---|---|---|---|---|---|---|---|---|---|
| | | Top 3 | Top 5 | 1% | 2% | 5% | 10% | 321.9 | 160.9 | 80.5 | 32.2 | 16.1 |
| **DyScore** | **0.866 (0.007)** | **43.7 (1.6)** | **39.9 (0.4)** | **32.1 (0.6)** | **21.5 (0.6)** | **11.0 (0.2)** | **6.5 (0.1)** | **0.619 (0.012)** | **0.535 (0.009)** | **0.495 (0.009)** | **0.515 (0.011)** | **0.569 (0.011)** |
| Comparison of DyScore and other SOTA Scoring Methods | | | | | | | | | | | | |
| Vina | 0.731 (0.002) | 14.6 (2.4) | 13.0 (2.1) | 10.7 (1.6) | 8.2 (0.8) | 5.2 (0.1) | 3.8 (0.1) | 0.208 (0.031) | 0.186 (0.025) | 0.185 (0.020) | 0.226 (0.013) | 0.291 (0.010) |
| DSX | 0.678 (0.004) | 11.2 (2.1) | 10.2 (1.5) | 8.0 (1.0) | 6.2 (0.5) | 4.2 (0.1) | 3.0 (0.1) | 0.152 (0.028) | 0.140 (0.018) | 0.142 (0.012) | 0.178 (0.010) | 0.233 (0.010) |
| RF-Score-v3 | 0.674 (0.005) | 7.3 (0.7) | 6.6 (0.3) | 5.2 (0.2) | 4.3 (0.3) | 3.4 (0.1) | 2.7 0.0 | 0.101 (0.006) | 0.093 (0.005) | 0.100 (0.005) | 0.139 (0.005) | 0.197 (0.003) |
| X-Score | 0.673 (0.003) | 6.3 (0.3) | 6.0 (0.6) | 5.2 (0.4) | 4.4 (0.3) | 3.4 (0.1) | 2.7 (0.1) | 0.099 (0.005) | 0.093 (0.007) | 0.101 (0.006) | 0.140 (0.005) | 0.198 (0.005) |
| NNScore-v2 | 0.624 (0.006) | 1.7 (0.6) | 2.0 (0.2) | 2.0 (0.6) | 1.9 (0.2) | 2.0 (0.3) | 1.8 (0.1) | 0.031 (0.011) | 0.035 (0.008) | 0.044 (0.007) | 0.076 (0.009) | 0.126 (0.010) |
| Score2 | 0.614 (0.001) | 1.2 (0.7) | 1.4 (0.1) | 1.8 (0.2) | 1.9 (0.1) | 2.1 (0.1) | 1.8 (0.1) | 0.026 (0.006) | 0.031 (0.004) | 0.043 (0.003) | 0.077 (0.003) | 0.128 (0.004) |
| Comparison of DyScore and other methods learning from the same features | | | | | | | | | | | | |
| DyScore-MLP | 0.849 (0.007) | 34.2 (1.1) | 32.2 (0.9) | 25.7 (0.5) | 18.1 (0.6) | 10.1 (0.4) | 6.1 (0.2) | 0.488 (0.014) | 0.432 (0.013) | 0.413 (0.015) | 0.451 (0.016) | 0.515 (0.016) |
| DyScore-RF | 0.854 (0.007) | 38.8 (2.1) | 34.5 (1.5) | 28.5 (0.7) | 19.7 (0.2) | 10.5 (0.3) | 6.4 (0.2) | 0.545 (0.017) | 0.478 (0.013) | 0.450 (0.008) | 0.482 (0.010) | 0.542 (0.013) |
| DyScore-DT-XGBoost | 0.859 (0.007) | 41.9 (1.7) | 38.2 (0.5) | 30.3 (0.5) | 20.6 (0.2) | 10.6 (0.1) | 6.3 (0.2) | 0.589 (0.013) | 0.510 (0.011) | 0.473 (0.009) | 0.496 (0.008) | 0.551 (0.009) |

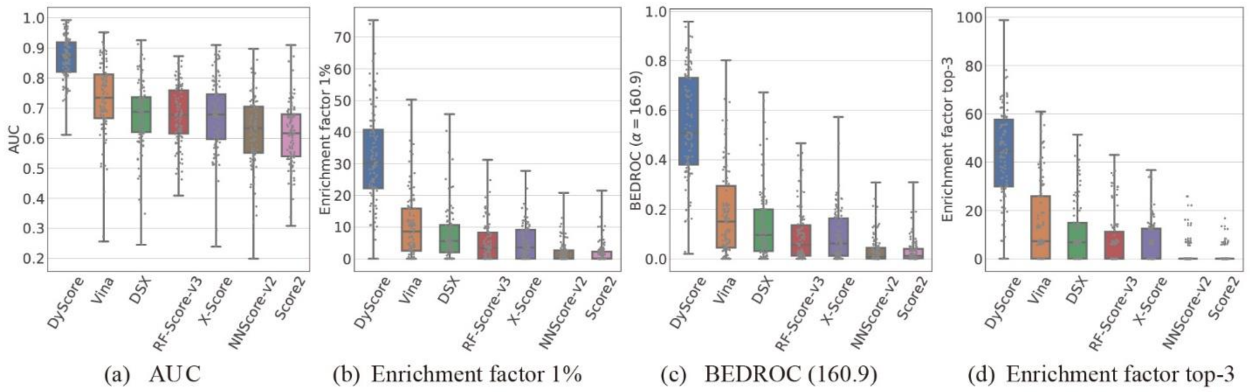


**Figure S3** Comparison results of different scoring methods on the DUD-E validation set generated by target-aware split. Each gray dot represents a target from DUD-E dataset (102 in total), and every point value is averaged over three random split validation sets.

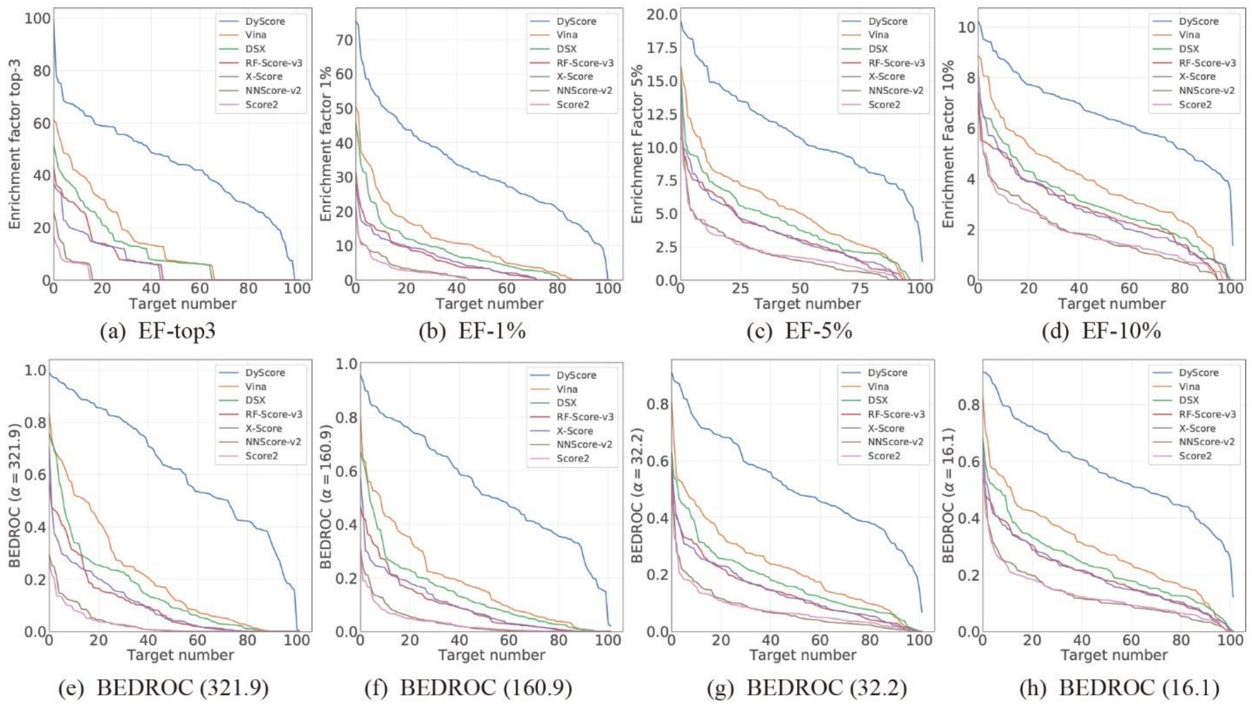


**Figure S4:** Enrichment factor and BEDROC to evaluate the performance of different scoring methods with respect to the target number. The metrics are measured on the DUD-E validation set generated by the target-aware split.

#### 1.4 Comparison on the DUD-E Validation Set with Target-unaware Split

**Table S2:** Comparison of DyScore with SOTA scoring functions on the DUD-E validation set generated by the target- unaware split. All the methods are evaluated on the three independent random split validation sets and the mean value is reported as well as the standard deviation in parentheses.

| Methods | AUC | Enrichment Factor | | | | | | BEDROC | | | | |
|---|---|---|---|---|---|---|---|---|---|---|---|---|
| | | Top 3 | Top 5 | 1% | 2% | 5% | 10% | 321.9 | 160.9 | 80.5 | 32.2 | 16.1 |
| **DyScore** | **0.866 (0.009)** | **44.2 (2.2)** | **38.6 (2.4)** | **30.8 (1.8)** | **21.2 (1.2)** | **11.0 (0.4)** | **6.5 (0.1)** | **0.632 (0.031)** | **0.541 (0.029)** | **0.496 (0.025)** | **0.515 (0.021)** | **0.568 (0.019)** |
| Vina | 0.727 (0.008) | 14.7 (1.1) | 12.7 (0.3) | 9.7 (0.2) | 7.3 (0.3) | 5.1 (0.2) | 3.7 (0.1) | 0.209 (0.009) | 0.180 (0.002) | 0.175 (0.003) | 0.215 (0.006) | 0.280 (0.008) |
| DSX | 0.673 (0.004) | 10.8 (1.3) | 9.9 (0.4) | 7.6 (0.8) | 6.3 (0.2) | 4.3 (0.2) | 3.1 (0.1) | 0.156 (0.015) | 0.142 (0.009) | 0.145 (0.004) | 0.183 (0.002) | 0.238 (0.001) |
| RF-Score-v3 | 0.668 (0.009) | 8.1 (0.5) | 7.3 (0.9) | 5.9 (0.6) | 4.7 (0.6) | 3.6 (0.2) | 2.7 (0.1) | 0.111 (0.007) | 0.104 (0.009) | 0.110 (0.009) | 0.148 (0.008) | 0.206 (0.007) |
| X-Score | 0.630 (0.005) | 6.8 (0.6) | 6.5 (0.1) | 5.6 (0.2) | 4.6 (0.1) | 3.5 0.0 | 2.7 (0.1) | 0.116 (0.010) | 0.105 (0.005) | 0.108 (0.004) | 0.144 (0.002) | 0.199 (0.001) |
| NNScore-v2 | 0.621 (0.003) | 1.9 (0.2) | 2.2 (0.2) | 2.1 0.0 | 2.1 0.0 | 1.9 (0.2) | 1.8 (0.1) | 0.033 (0.003) | 0.037 0.000 | 0.047 (0.003) | 0.079 (0.005) | 0.129 (0.004) |
| Score2 | 0.609 (0.007) | 2.4 (1.2) | 2.2 (0.4) | 2.2 (0.3) | 2.0 (0.2) | 2.0 (0.2) | 1.9 (0.1) | 0.035 (0.008) | 0.038 (0.004) | 0.047 (0.004) | 0.078 (0.006) | 0.128 (0.006) |

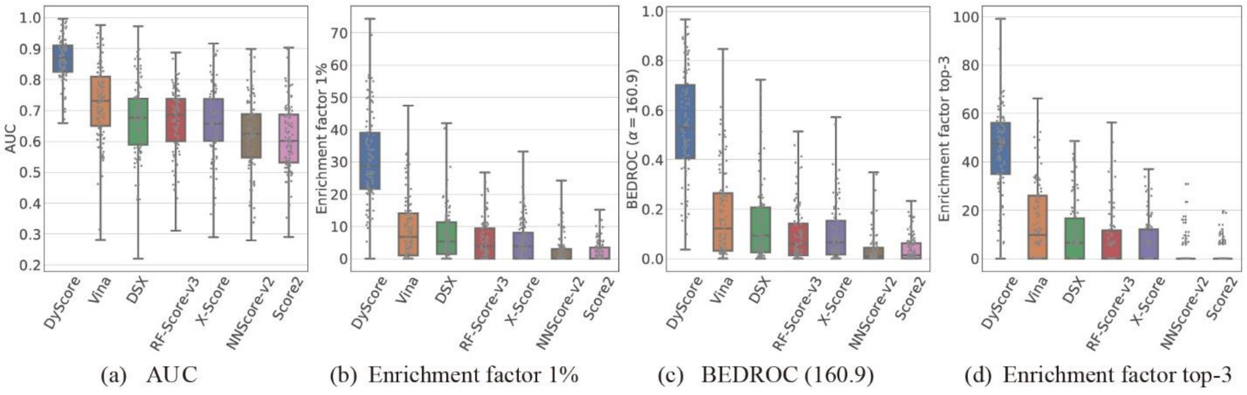


**Figure S5:** Comparison results of different scoring methods on the DUD-E validation set generated by target-unaware split. Each gray dot represents a target.

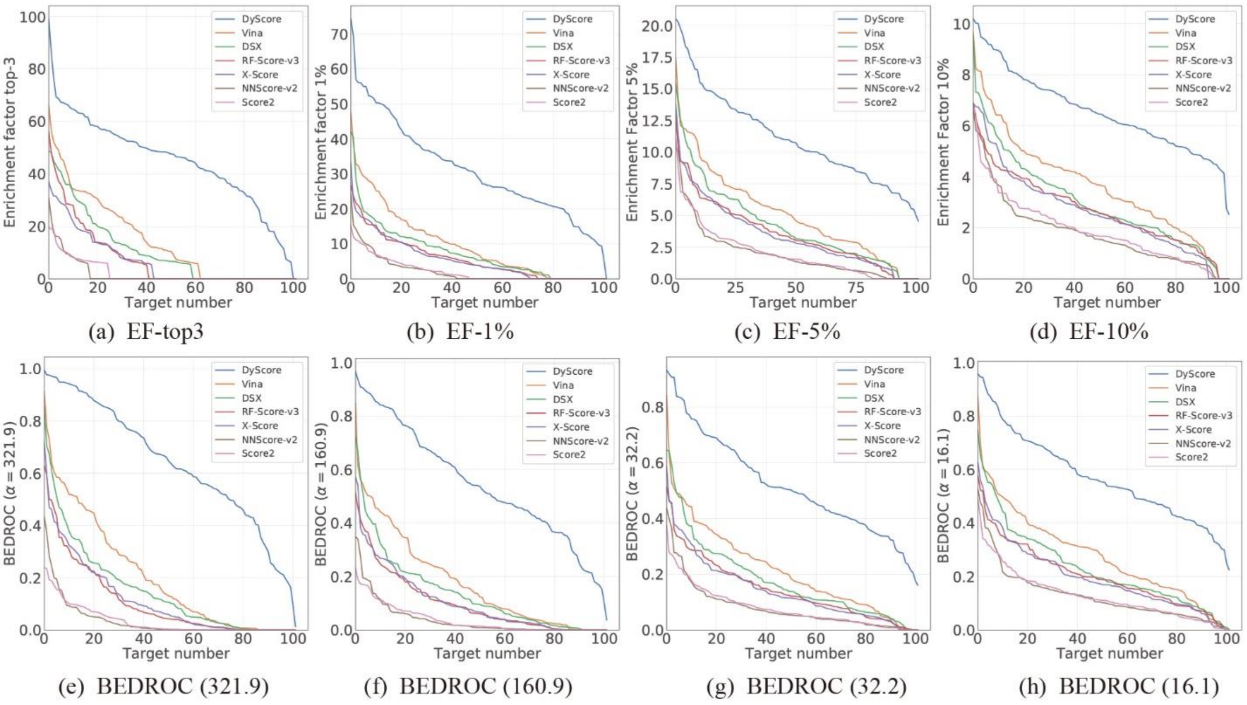


**Figure S6:** Enrichment factor and BEDROC to evaluate the performance of different scoring methods with respect to the target number. The metrics are measured on the DUD-E validation set generated by the target-unaware split.

#### 1.5 Per-target ROC curves of DyScore for the DUD-E test set with target-aware split

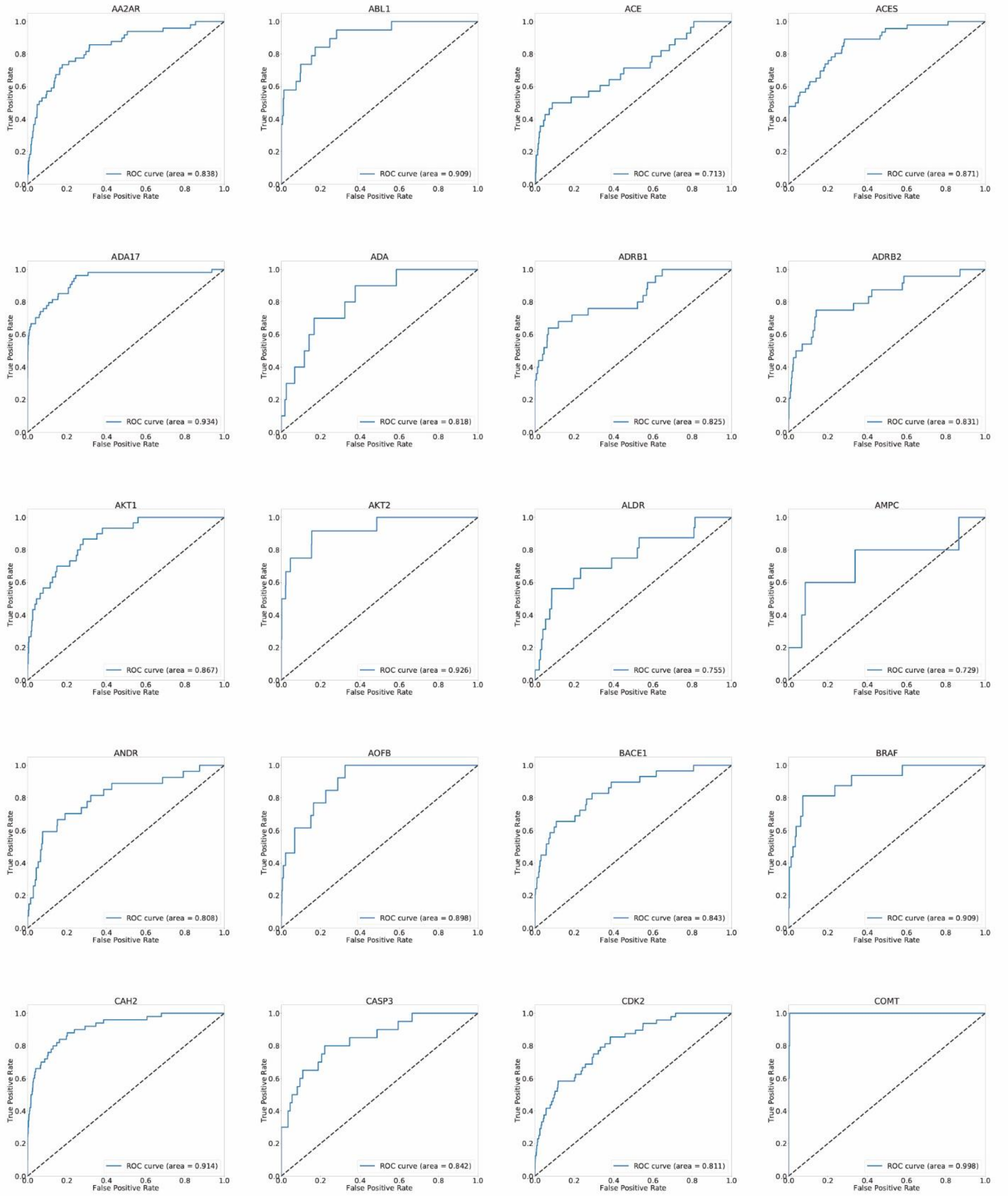


**Figure S7:** ROC curves for each target in DUD-E dataset. (Cont.)

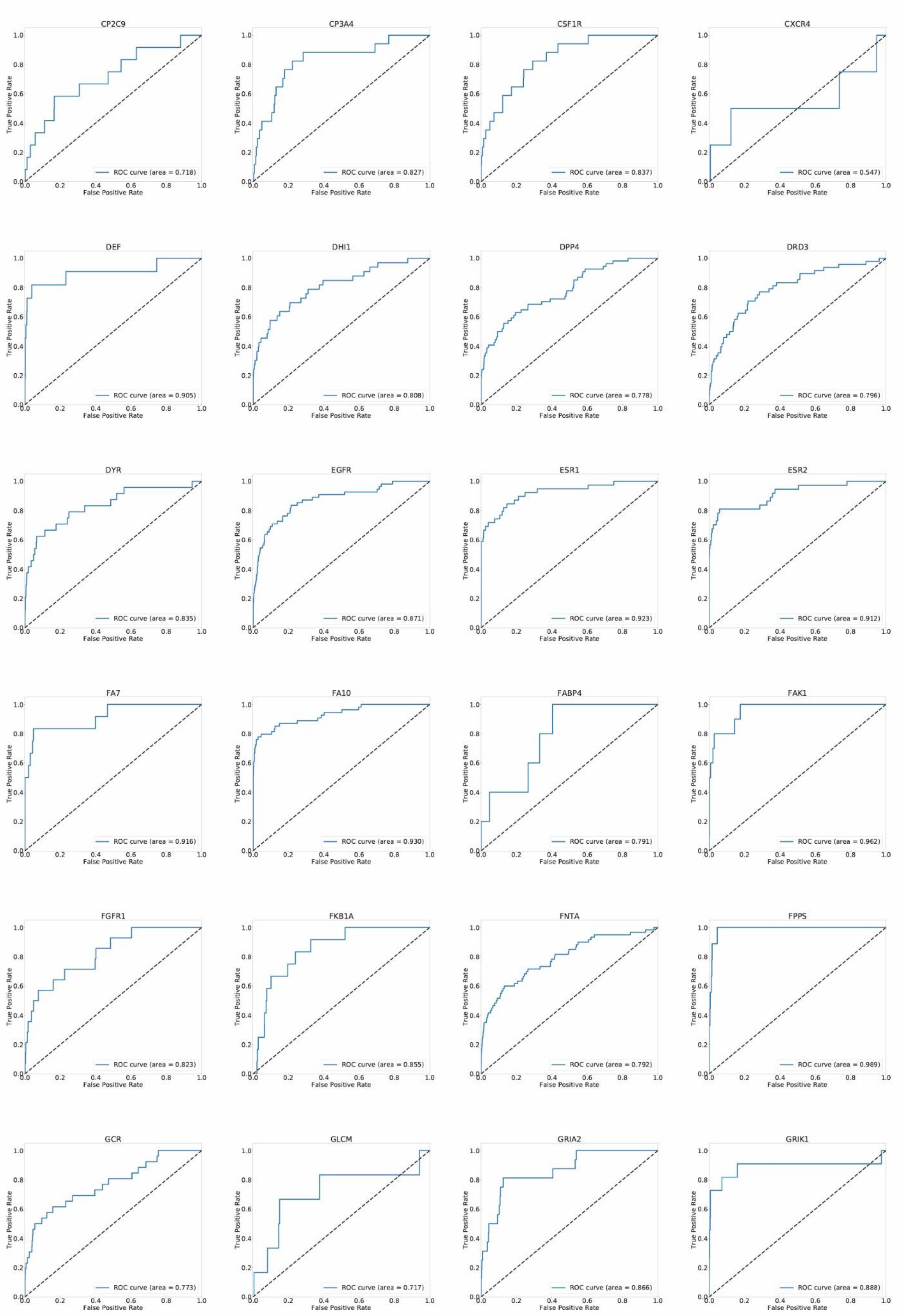


**Figure S7:** ROC curves for each target in DUD-E dataset (Cont.).

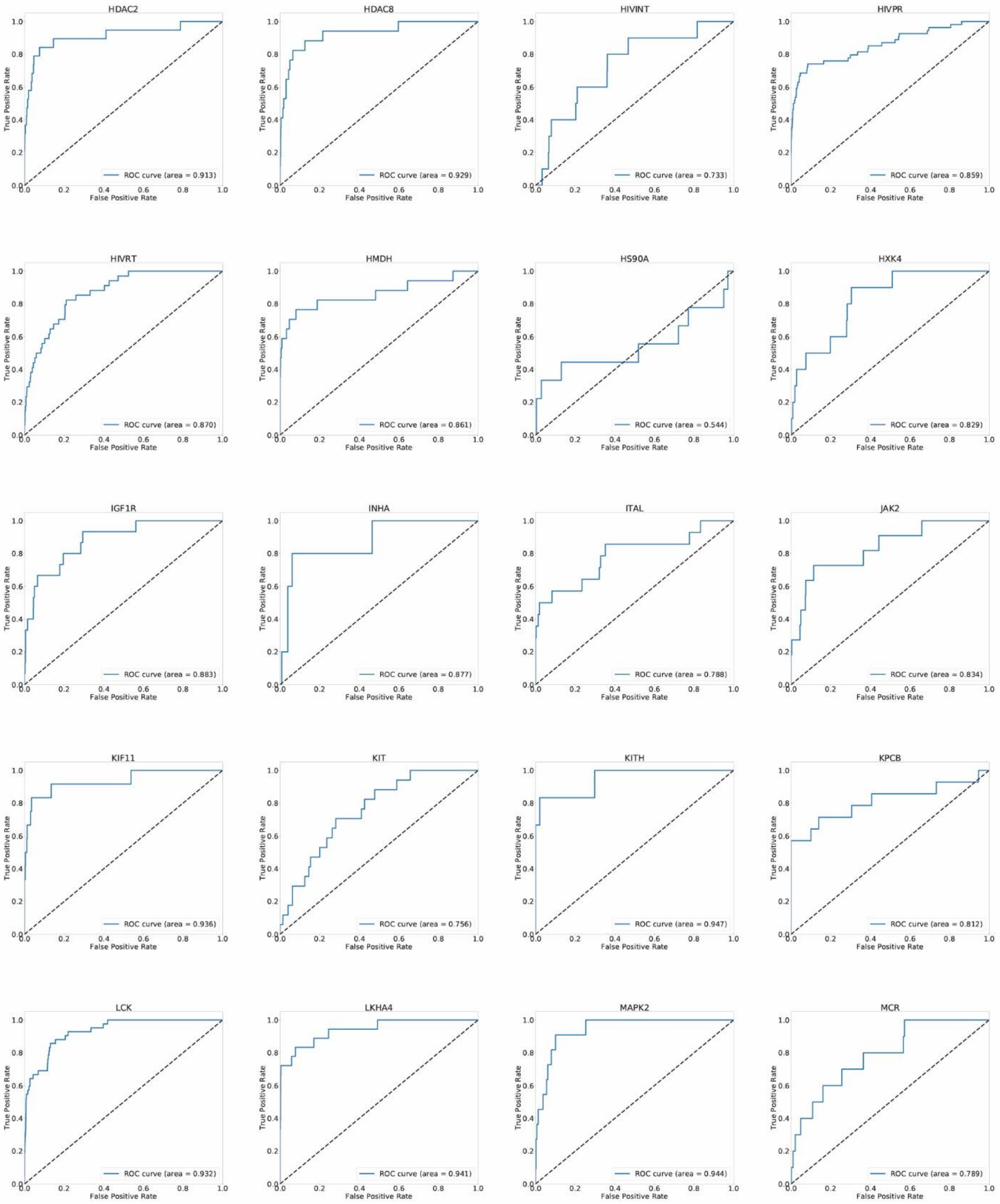


**Figure S7:** ROC curves for each target in DUD-E dataset (Cont.).

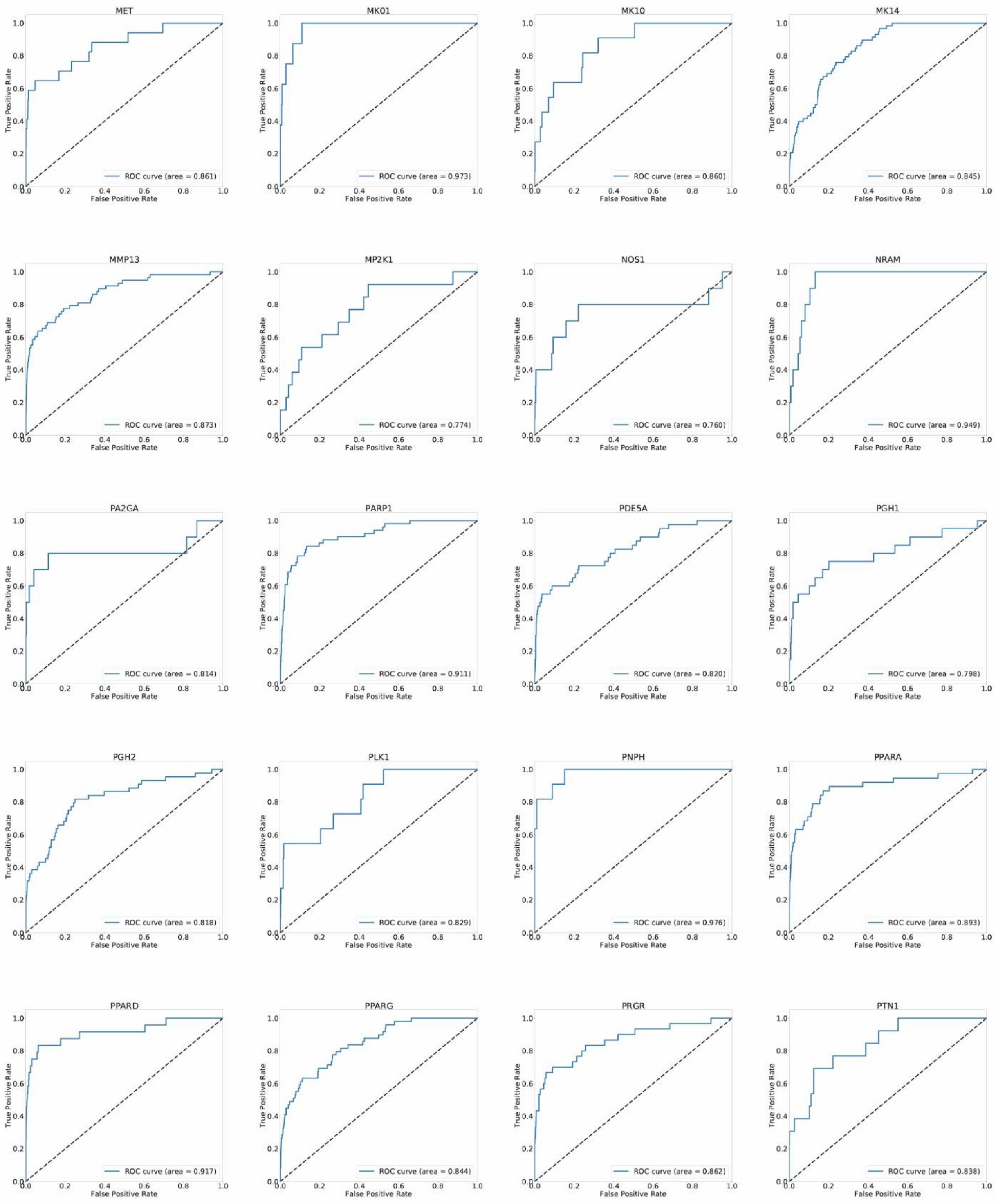


**Figure S7:** ROC curves for each target in DUD-E dataset (Cont.).

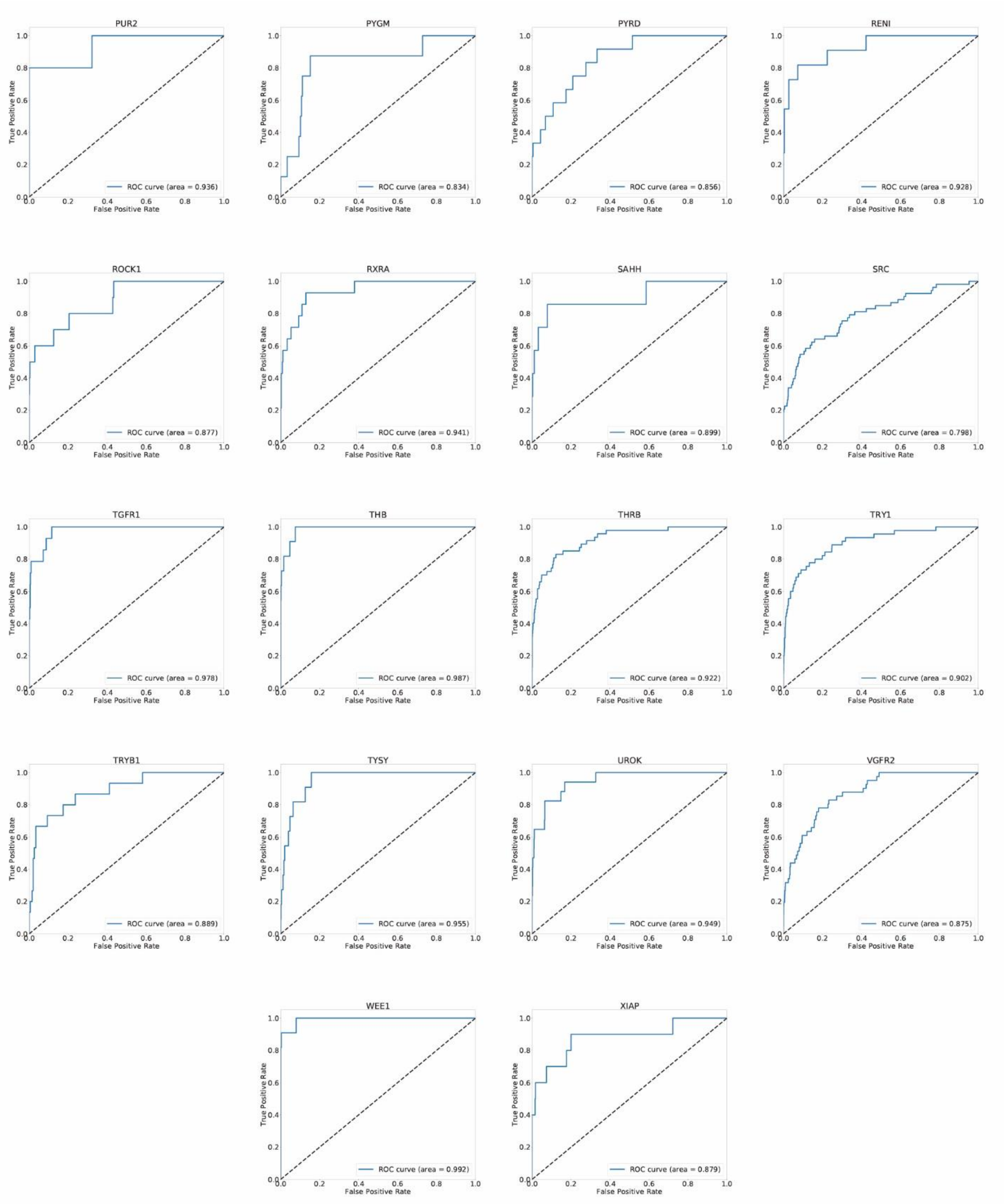


**Figure S7:** ROC curves for each target in DUD-E dataset.

### 1.6 Per-target ROC curves of DyScore for the DEKOIS 2.0 data set

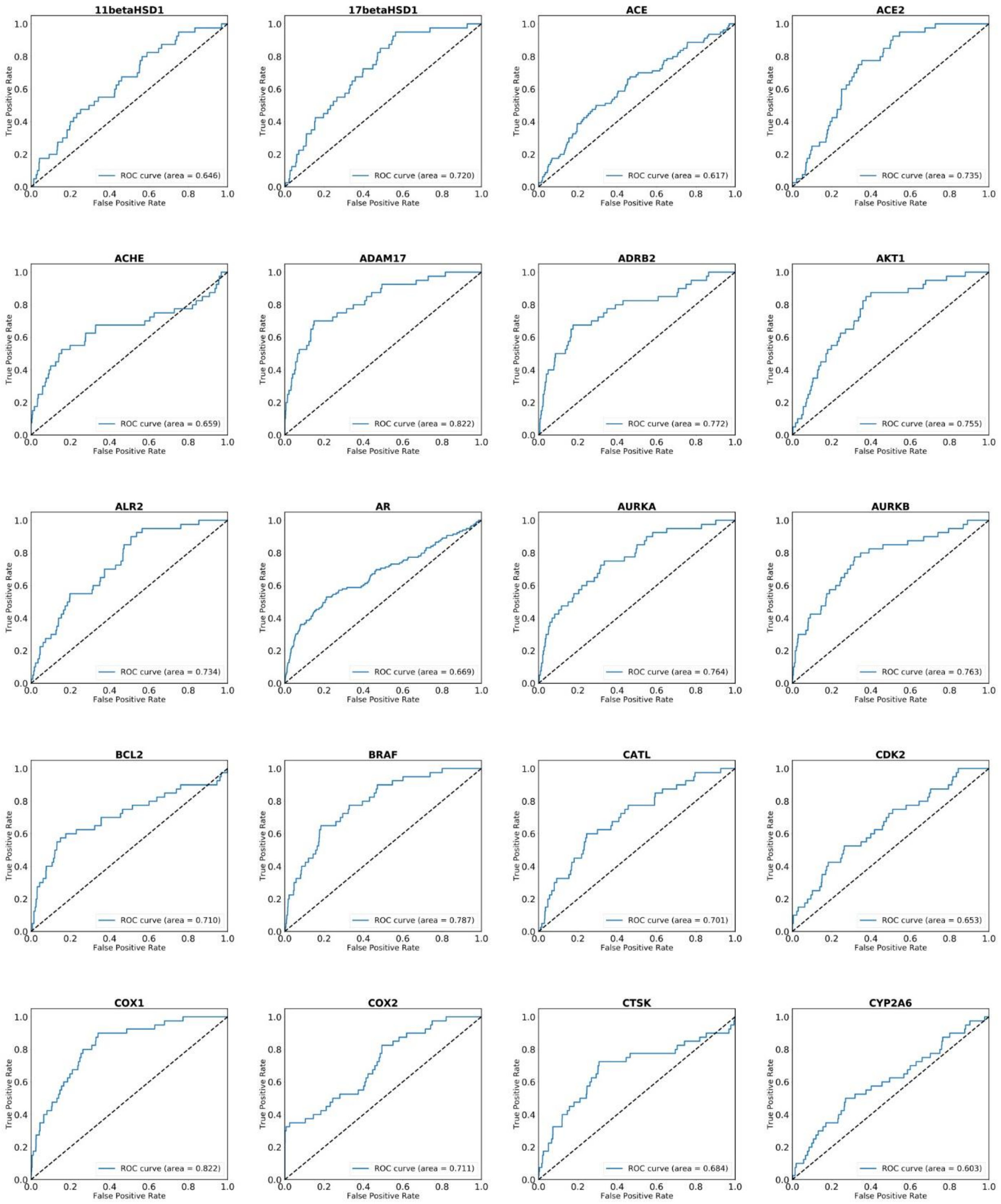


**Figure S8:** ROC curves for each target in DEKOIS 2.0 benchmark. (Cont.)

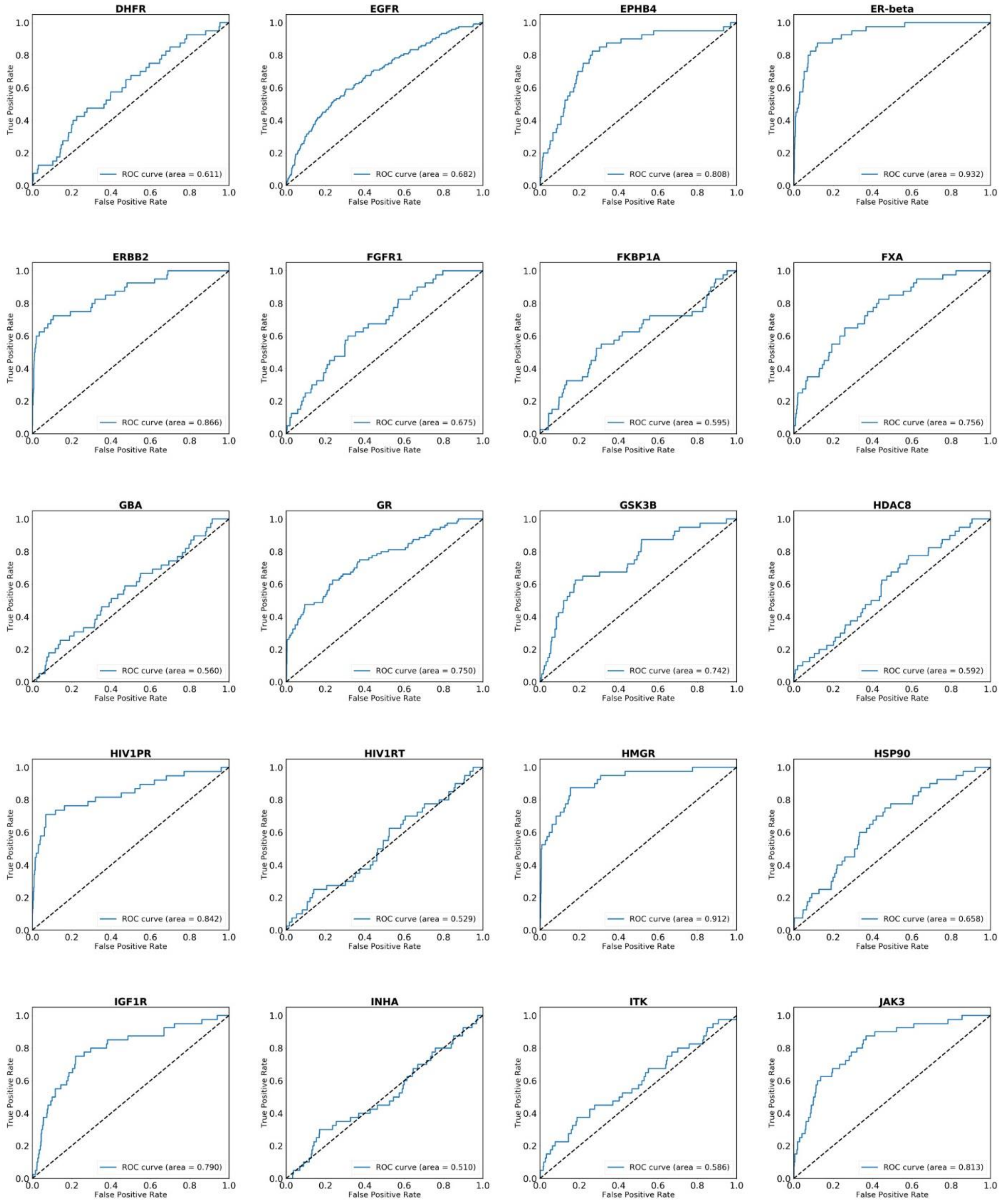


**Figure S8:** ROC curves for each target in DEKOIS 2.0 benchmark. (Cont.)

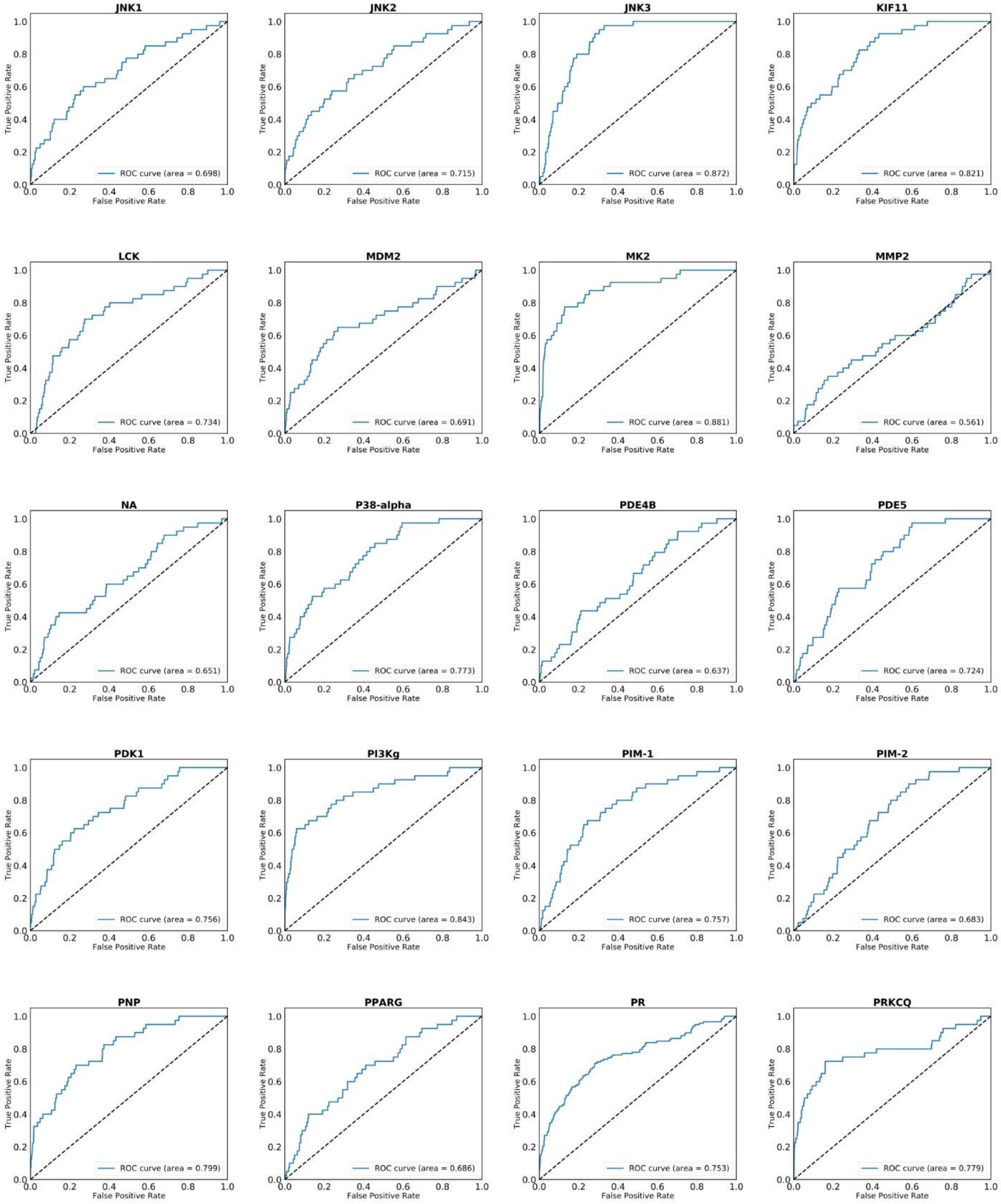


**Figure S8:** ROC curves for each target in DEKOIS 2.0 benchmark (Cont.)

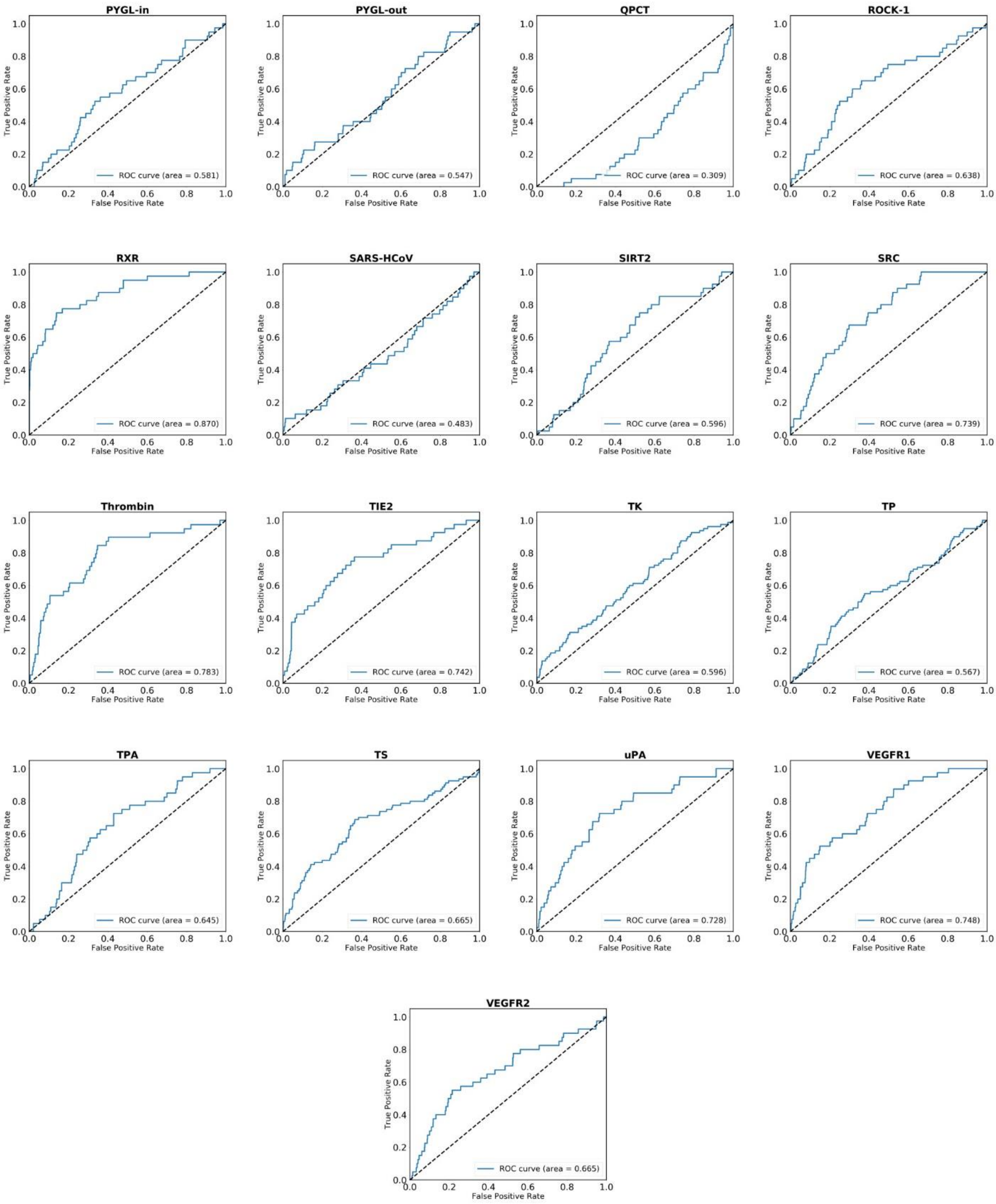


**Figure S8:** ROC curves for each target in DEKOIS 2.0 benchmark.

#### 1.6 Comparison of DyScore and Dyscore-MF with SOTA scoring function on LIT-PCBA database.

**Table S3:** Comparison of DyScore and DyScore-MF with six baseline SOTA scoring functions on the LIT-PCBA dataset.

| Methods | AUC | Enrichment Factor | | | | | | BEDROC | | | | |
|---|---|---|---|---|---|---|---|---|---|---|---|---|
| | | Top 3 | Top 5 | 1% | 2% | 5% | 10% | 321.9 | 160.9 | 80.5 | 32.2 | 16.1 |
| DyScore-MF | 0.594 | 9.5 | **11.7** | **5.9** | **4.7** | **3.0** | **2.3** | **0.043** | **0.054** | **0.071** | **0.110** | **0.161** |
| DyScore | 0.563 | **11.1** | 6.7 | 3.3 | 2.7 | 2.2 | 1.7 | 0.024 | 0.030 | 0.043 | 0.075 | 0.117 |
| Vina | 0.565 | 1.1 | 2.1 | 2.3 | 1.9 | 1.7 | 1.5 | 0.030 | 0.030 | 0.037 | 0.063 | 0.104 |
| RF-Score-v3 | 0.571 | 0.0 | 0.0 | 2.0 | 1.6 | 1.7 | 1.4 | 0.017 | 0.022 | 0.031 | 0.058 | 0.100 |
| NNScore-v2 | 0.557 | 0.0 | 0.0 | 1.7 | 1.4 | 1.5 | 1.3 | 0.008 | 0.014 | 0.025 | 0.052 | 0.092 |
| DSX | 0.523 | 0.0 | 0.7 | 1.0 | 1.7 | 1.4 | 1.2 | 0.014 | 0.017 | 0.025 | 0.049 | 0.085 |
| Score2 | **0.621** | 0.0 | 8.3 | 3.0 | 3.1 | 2.2 | 1.9 | 0.023 | 0.032 | 0.047 | 0.081 | 0.131 |
| Xscore | 0.576 | 0.0 | 0.7 | 2.6 | 2.5 | 1.9 | 1.5 | 0.022 | 0.029 | 0.039 | 0.067 | 0.110 |

**Table S4:** Comparison of EF1% for DyScore and DyScore-MF with twelve SOTA methods on the LIT-PCBA dataset.

| **Methods** | **Enrichment Factor 1% (EF0.01)** | **Ref** |
|---|---|---|
| RFScore-4 | 1.28 | Sunseri et al. [1] |
| RFScore-VS | 0.73 | |
| GNINA | 2.58 | |
| 2D ECFP4 similarity search* | 2.49 | Zhang et al. [2] |
| 3D shape similarity search* | 0.96 | |
| EVIS-best | 4.18 | |
| EVIS-ave* | 2.31 | |
| FINDSITE$^{comb2.0}$ | 3.04 | Zhou et al. [3] |
| FRAGSITE | 4.78 | |
| Vina | 2.333 | This study |
| RF-Score-v3 | 1.999 | |
| NNScore-v2 | 1.679 | |
| DSX | 1.012 | |
| Score2 | 2.954 | |
| Xscore | 2.583 | |
| Dyscore | 3.34 | |
| **DyScore-MF** | **5.92** | |

*For reference, not counted in the twelve SOTA methods.

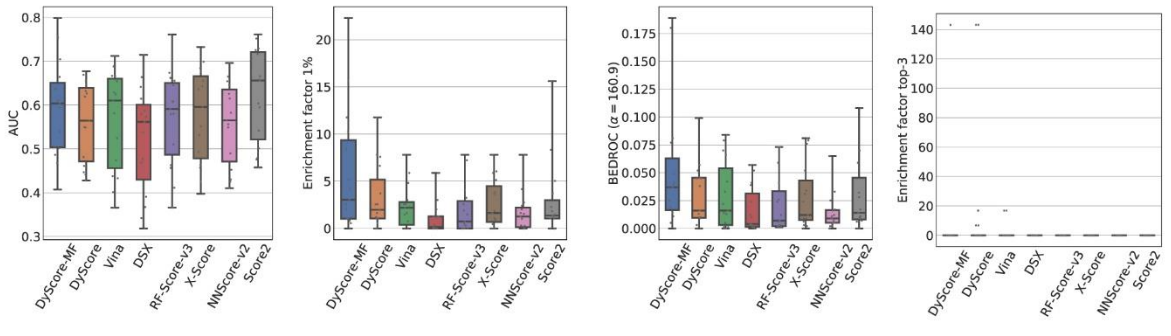


(a) AUC (b) Enrichment factor 1% (c) BEDROC (160.9) (d) Enrichment factor top-3

**Figure S9:** Comparison results of different scoring methods on the LIT-PCBA dataset.

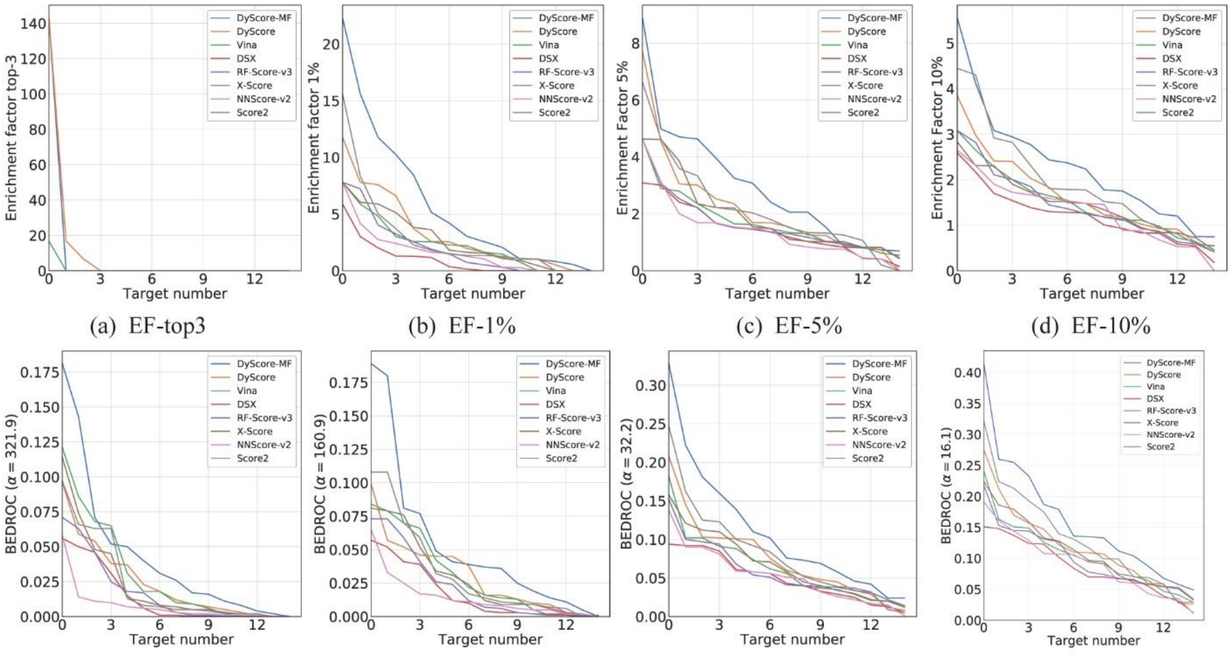


(a) EF-top3 (b) EF-1% (c) EF-5% (d) EF-10%

(e) BEDROC (321.9) (f) BEDROC (160.9) (g) BEDROC (32.2) (h) BEDROC (16.1)

**Figure S10:** Enrichment factor and BEDROC to evaluate the performance of different scoring methods with respect to the target number on LIT-PCBA dataset.

#### 1.7 Computational cost DyScore and DyScore-MF

**Table S5:** Statistic of computational cost per thousand compounds per CPU in each step. Tested with AMD EPYC 75F3 Milan.

| Step | Time (second/1K compounds /CPU) |
|---|---|
| RF-Score-v3 | 331 |
| DSX | 164 |
| Score2 | 180 |
| Xscore | 794 |
| **NNScore2 (Vina score)*** | **2552** |
| Fingerprint & other features | 151 |
| **Dynamic Features** | **913** |
| Model Prediction | 83 |
| Total | 5168 |

*Only calculate the binding score via autodock vina, instead of performing the molecular docking.
